# Early sexual dimorphism in the developing gut microbiome of northern elephant seals

**DOI:** 10.1101/2019.12.11.871855

**Authors:** M.A. Stoffel, K. Acevedo-Whitehouse, Nami Morales-Durán, S. Grosser, N. Chakarov., O. Krüger, H.J. Nichols, F.R. Elorriaga-Verplancken, J.I. Hoffman

## Abstract

The gut microbiome is an integral part of a species’ ecology, but we know little about how host characteristics impact its development in wild populations. Here, we explored the role of such intrinsic factors in shaping the gut microbiome of northern elephant seals during a critical developmental window of six weeks after weaning, when the pups stay ashore without feeding. We show that the early-life gut microbiome is already substantially different in male and female pups, even though males and females cannot yet be distinguished morphologically. Sex and age both explain around 15% of the variation in gut microbial beta diversity, while microbial communities sampled from the same individual show high levels of similarity across time, explaining another 40% of the variation. Only a small proportion of the variation in beta diversity is explained by health status, but healthy individuals have a greater microbial alpha diversity than their non-healthy peers. Across the post-weaning period, the elephant seal gut microbiome is highly dynamic. We found evidence for several colonisation and extinction events as well as a decline in *Bacteriodes* and an increase in *Prevotella*, a pattern that has previously been associated with the transition from nursing to solid food. Lastly, we show that genetic relatedness is correlated with gut microbiome similarity in males but not females, again reflecting substantial early sex-differences. Our study represents a naturally diet-controlled and longitudinal investigation of how intrinsic factors shape the early gut microbiome in a species with extreme sex differences in morphology and life history.

## Introduction

Vertebrates are inhabited by vast numbers of microbes that are increasingly emerging as key players in their host’s biology and evolution (Bik et al., 2016; Ley et al., 2008; McFall-Ngai et al., 2013; Moeller et al., 2014). The richest and arguably most complex microbial communities are those that populate the gastrointestinal tract and which are collectively termed the ‘gut microbiome’. Gut microbes benefit their host in many ways, such as promoting the development of organs, assisting nutrient uptake and priming and modulating the immune system (Cheesman, Neal, Mittge, Seredick, & Guillemin, 2011; Heijtz et al., 2011; Lathrop et al., 2011; Zhu, Wu, Dai, Zhang, & Wei, 2011). Consequently, disturbances to the gut microbiome can have severe consequences for the host, ranging from autoimmune diseases and infections to obesity (Giongo et al., 2011; Round & Mazmanian, 2009; Turnbaugh, Bäckhed, Fulton, & Gordon, 2008).

The gut microbiome is highly dynamic across space and time and can be influenced by many factors. On a broader scale, the strongest determinants of the gut microbiome appear to be phylogeny and diet, both of which can result in remarkably different bacterial communities across host species (Bik et al., 2016; Ley et al., 2008; Muegge et al., 2011). On a finer scale, differences in the gut microbiome within species can be shaped by a combination of environmental factors such as diet, location and season, behavioural factors such as social networks, and heritable factors such as host genetics (Benson et al., 2010; Kurilshikov, Wijmenga, Fu, & Zhernakova, 2017; Moeller et al., 2014; Ren et al., 2017; Tung et al., 2015). However, most studies to date have focused on animals held in captivity, which can influence microbial communities due to factors such as controlled and less diverse diets (Hird, 2017). Consequently, relatively little is known about the composition, development and function of the gut microbiome in the wild, despite its potential to contribute to our fundamental understanding of the ecology and evolutionary biology of mutualistic symbiotic relationships (Hird, 2017; Zilber-Rosenberg & Rosenberg, 2008).

The mammalian foetal gut is considered to be largely sterile, although there is recent evidence of uterine bacterial translocation to the foetus (Chen & Gur, 2019; Perez-Muñoz, Arrieta, Ramer-Tait, & Walter, 2017). During and after birth, the gut becomes rapidly colonised by various microbes. In these early stages of life, the gut microbiome is of tremendous importance and disturbances can impact host development and impair metabolism, health and immune function (Candon et al., 2015; Cho et al., 2012; Cox et al., 2014; Macpherson & Harris, 2004; Russell et al., 2012). It is therefore of interest to investigate changes in the microorganisms that populate the gut during an individual’s development. Across the life-span of an organism, ontogeny appears to influence the composition of the gut microbiome in a species-specific manner (Clark et al., 2015; Langille et al., 2014; O’Toole & Jeffery, 2015). For example, bacterial diversity increases throughout early development in humans, chickens, pigs and ostriches (Ballou et al., 2016; Frese, Parker, Calvert, & Mills, 2015; Kundu, Blacher, Elinav, & Pettersson, 2017; Videvall et al., 2019), but decreases during maturation in zebrafish and African turquoise killifish (Smith et al., 2017; Stephens et al., 2016). A mixed pattern has been observed in mice, where an early drop in diversity after the initial transmission of maternal microbiota is followed by an increase after the introduction of solid food (Pantoja-Feliciano et al., 2013). However, to our knowledge, patterns of microbial colonisation during early development in wild animals are as yet largely unknown (Ren et al., 2017).

Every species’ life-history is characterised by a series of challenges to which an organism must adapt, both physiologically and behaviourally. A key element facilitating these adaptations might be the microbiome. A particularly strong factor driving within-species variation in microbial communities could be sex, as males and females often experience contrasting selection pressures due to differences in their behaviour and physiology (Tarka, Guenther, Niemelä, Nakagawa, & Noble, 2018). Several of these differences might be directly or indirectly associated with the gut microbiome, such as sex-specific immune responses (Klein & Flanagan, 2016) or sex-specific foraging behaviour (Boeuf et al., 2000; Boinski, 1988; Lewis et al., 2002). Surprisingly given the important role of sex-specific microbiota in humans (Markle et al., 2013), the impact of sex on the gut microbiome of wild vertebrates seems to be non-existent or very small (Bennett et al., 2016; Bobbie, Mykytczuk, & Schulte-Hostedde, 2017; Maurice et al., 2015; Ren et al., 2017; Tung et al., 2015). However, gut microbiome studies of wild populations are likely to be impacted by environmental factors that can rarely be controlled for and which could potentially mask any effects of intrinsic factors such as sex.

Another largely open question for natural populations is how host genotype affects the gut microbiome. Most insights to date come from twin studies in humans or from different strains of laboratory mice and suggest that the influence of host genetics is modest compared to environmental effects (Kurilshikov et al., 2017). However, most free-ranging animal populations carry greater levels of genetic variation than inbred laboratory stocks and their microbiota may also be more complex, which could potentially lead to stronger covariation between host genotype and microbial community composition. Nevertheless, quantifying the impact of host genetics on the gut microbiome in the wild is challenging, at least in part because of the need to control for environmental effects (Bik et al., 2016; Perofsky, Lewis, Abondano, Di Fiore, & Meyers, 2017; Tung et al., 2015) that may might blur any genetic signal. Consequently, it remains unclear whether host genetics influences the gut microbiome in natural populations, despite the importance of this question in the light of host-microbe evolution.

An ideal opportunity to investigate the intrinsic factors that shape the gut microbiome in the wild is provided by the northern elephant seal (*Mirounga angustirostris*). Northern elephant seals are among the most sexually dimorphic of all mammals, with males being 3-4 times heavier than females (Wilson & Mittermeier, 2014). The mating system of this species is highly polygynous, with only a handful of successful males copulating with dozens of females in a given season (Burney J. Le Boeuf & Laws, 1994). Consequently, males and females face very different challenges: during the breeding season, males must continuously defend their harems against competitors, while females need to invest substantial amounts of energy into nursing their pups. Neither males nor females feed during the breeding season, with some males fasting for up to three months and females fasting for up to one month, despite the high energetic investment required to provide high-fat milk to their young (Burney J. Le Boeuf & Ortiz, 1977). Outside the breeding season, elephant seals spend most of their lives at sea, and even there, sex differences are apparent. Adult males and females have very different foraging strategies, with males feeding on benthic prey along the continental margin of North America, and females feeding largely on pelagic prey in deeper waters (Boeuf et al., 2000). Consequently, elephant seals have developed a series of sex-specific adaptations to these diverging life-histories, but it is not known yet whether or how the gut microbiome might be involved.

Here, we studied the gut microbiome of elephant seal pups over a 35-day post-weaning period commencing immediately after their mothers stop nursing. This time-window is ideally suited to investigating the influence of intrinsic factors on gut microbiomes because all northern elephant seal pups remain within their natal colonies without feeding until they leave the rookery around seven weeks later (Reiter, Stinson, & Boeuf, 1978). Consequently, variation in gut microbiome beta diversity (microbiome similarity between samples) and alpha diversity (microbiome diversity within samples) should be largely driven by intrinsic factors such as sex, developmental stage and health rather than by extrinsic factors such as habitat or dietary changes. We therefore used repeated, longitudinal sampling of rectal swabs to characterise the early-life gut microbiome of the northern elephant seal and to explore the factors driving variation in beta and alpha diversity, with a particular emphasis on sex-differences, which may reflect early life-history adaptations. Lastly, we used microsatellite genotyping to test the hypothesis that genetically more related individuals also host more similar gut microbiomes. Overall, our study provides a rare glimpse into the early development of the gut microbiome in a natural population within a diet-controlled setting that allowed us to evaluate intrinsic sources of microbial variation in the wild.

## Materials and Methods

### Study design and sample collection

We marked 40 northern elephant seal pups and their mothers during the breeding season in February/March 2017 at Benito del Oeste, the westernmost island of the San Benito Archipelago off the west coast of Baja California, Mexico. We closely observed mother-offspring pairs in order to determine the weaning dates of each pup. Weaning typically occurs close to 28 days after birth (Reiter et al., 1978) and marks the time that the mother abandons her pup and returns to the sea. At this moment, we sampled the newly weaned pup (time point T1). To analyse the gut microbiome composition, we took rectal swabs using FLOQSwabs™, which were immediately stored in 70% EtOH, frozen at −20°C within a few hours after collection and subsequently stored at −80°C shortly after the end of the field season. To determine the genetic relatedness of individuals, we collected a small skin sample (9 mm^2^) from the flipper of each pup and stored these individually in sterile cryogenic vials containing 70% EtOH. The vials were frozen at −20°C within a few hours after collection and were subsequently stored at −80°C. During the T1 sampling period, we collected rectal swabs and skin samples from 40 pups, which were marked with plastic flipper tags with a unique ID number. Subsequently, we observed these pups on a daily basis and captured them after 15 days (T2) and 30 days (T3) to collect two additional rectal swabs for microbial profiling. The entire sampling scheme spanned the two-month long fasting period during which the weaned pups stay ashore (Reiter et al., 1978). We sampled blood from each individual to assess its health status at T1 and T3, Briefly, blood was collected from the extradural intervertebral vein, using a vacuum blood collection device with a 18 G needle. Blood samples were preserved with EDTA and were used to determine the total and differential leukocyte counts as has been described previously (Flores-Morán et al., 2017). Throughout the field season, we lost six of the marked pups, as one died between T1 and T2, one between T2 and T3, one was not found after T1 and three pups were lost after T2, despite intensive searching effort. Thus, sample sizes were 40 pups at T1, 38 at T2, and 34 at T3. All sampling was conducted with the approval of the Bioethics Committee and IACUC of the Autonomous University of Queretaro, and all capture and sampling procedures were carried out under permit DGVS 00091 /17 issued by the Mexican Secretariat of the Environment and Natural Resources.

### Host DNA extraction and microsatellite genotyping

Total genomic DNA was extracted from skin samples using a standard chloroform extraction protocol and genotyped at 21 previously developed microsatellite loci (see Supplementary Information for details). We tested all of the microsatellite loci for deviations from Hardy-Weinberg equilibrium (HWE) using exact tests based on Monte Carlo simulations implemented in pegas (Paradis, 2010) and applied a false discovery rate correction (Benjamini & Hochberg, 1995) to the resulting *p*-values. All 21 loci were retained in the final dataset as no locus was out of HWE.

### Bacterial DNA extraction, library preparation and sequencing

We extracted DNA from 112 rectal swabs using the QIAamp PowerFecal DNA Kit (Qiagen), and amplified a 300 bp of the V3 and V4 regions of the 16S rRNA gene. The amplicon libraries were prepared as follows: 1-10 ng of DNA extract (total volume 1μl), 15 pmol of each forward primer 341F 5’-NNNNNNNNNNTCCTACGGGNGGCWGCAG and reverse primer 785R 5’-NNNNNNNNNNTGACTACHVGGGTATCTAAKCC in 20 μL volume of 1 x MyTaq buffer containing 1.5 units MyTaq DNA polymerase (Bioline) and 2 μl of BioStabII PCR Enhancer (Sigma). For each sample, the forward and reverse primers had the same 10-nt barcode sequence. PCRs were carried out for 30 cycles of 1 min 96°C pre-denaturation; 96°C for 15 s, 50°C for 30 s, 70°C for 90 s. The DNA concentration of the amplicons of interest was determined by gel electrophoresis. About 20 ng of amplicon DNA of each sample was pooled for up to 48 samples carrying different barcodes. The amplicon pools were purified with one volume of AMPure XP beads (Agencourt) to remove primer dimer and other small miss-priming products, followed by an additional purification on MiniElute columns (Qiagen). About 100 ng of each purified amplicon pool DNA was used to construct Illumina libraries using the Ovation Rapid DR Multiplex System 1-96 (NuGEN). Illumina libraries were pooled and size selected by preparative gel electrophoresis. Sequencing was performed on an Illumina MiSeq platform using V3 Chemistry – 2×300 bp read length (Illumina). DNA extraction, library preparation and sequencing were carried out by LGC Genomics in Berlin.

### Bioinformatics pipeline

The 16S sequences in FASTQ format were demultiplexed using the Illumina bcl2fastq 2.16.1.14 software while allowing up to 2 mismatches or Ns in the barcode. Reads were sorted according to their barcodes, allowing up to 1 mismatch per barcode before removing the barcodes. Reads with missing, one-sided or conflicting barcode pairs were discarded. Adapters were clipped using cutadapt 1.13 (Martin, 2011) and all reads smaller than 100 bp were filtered out. Amplicon primers were detected while allowing for up to three mismatches, and primer pairs (Forward-Reverse or Reverse-Forward) had to be present in the sequence fragments. If primer dimers were detected, the outer primer copies were clipped from the sequence and the sequence fragments were turned into forward-reverse primer orientation after removing the primer sequence.

We used DADA2 1.8 (Benjamin J. Callahan et al., 2016) for further filtering and processing the sequences into Amplicon sequence variants (ASVs), following the authors’ published workflow (Ben J. Callahan, Sankaran, Fukuyama, McMurdie, & Holmes, 2016). Unlike the traditional grouping into operational taxonomic units (OTUs), ASVs are exact sequence variants and have the compelling advantages of higher taxonomic resolution as well as reproducibility and reusability across studies (Benjamin J. Callahan, McMurdie, & Holmes, 2017). After visually inspecting the quality profiles of all reads, we used DADA2’s filterAndTrim function to trim R1 and R2 sequences to 220 and 230 base pairs respectively and to filter all reads with more than two expected errors (Edgar & Flyvbjerg, 2015). As DADA2 relies on a parametric error model, we used the learnErrors function to evaluate error rates from the data and visually confirmed that the resulting error rate estimates provided a good fit to the observed rates using plotErrors (Ben J. Callahan et al., 2016). After dereplication with derepFastq, we used the dada function for correcting substitution and indel errors as well as for sample inference based on the pooled samples. Subsequently, we merged forward and reverse reads with a minimum overlap of 12 bp using mergePairs and constructed a sequence table with makeSequenceTable. After inspecting the distribution of sequence lengths across samples and considering a median full amplicon size of around 460 bp prior to primer clipping (Klindworth et al., 2013), primer-clipped sequences of lengths between 380 and 450 bp were retained. As a last filtering step, we removed chimeras with removeBimeraDenovo using the consensus method. We assigned taxa to the ASVs using the assignTaxonomy and addSpecies functions based on the SILVA database v128 (Quast et al., 2012). The resulting ASV table contained 2809 ASVs in 112 samples.

### Clinical assessment of health

For each blood smear, we quantified the differential white blood cell populations by counting the number of lymphocytes, neutrophils, band neutrophils, hypersegmented neutrophils, monocytes, basophils, and eosinophils, in 100 leukocytes. Absolute numbers for each leucocyte type were calculated by multiplying the total white blood cell count, previously determined by the use of a Neubauer chamber, by the proportion of each leucocyte type. Based on the clinical reference values previously reported for clinically healthy northern elephant seal pups (Bossart, Reidarson, Dierauf, & Duffield, 2001; Yochem, Stewart, Mazet, & Boyce, 2008) we classified the clinical health status of each pup as either healthy (i.e. none of the leukocyte types deviated from the normal ranges) or not-healthy (i.e. at least one cell type was out of the normal range).

### Data processing and analyses

#### Microbial data

All subsequent analyses were conducted in R version 3.4.3 (R Core Team). As a first filtering step after taxonomic assignment, we discarded ASVs classified as mitochondria (*n* = 3) or chloroplasts (*n* = 8) together with ASVs that could not be identified at the Class level (*n* = 77), as these are more likely to contain sequencing errors. Based on a visual assessment of ASV abundance and prevalence (Supplementary Figure 5), we then removed ASVs that did not appear in at least three samples (*n* = 982) or which had a total read count below 30 across all samples (*n* = 683).

Overall, 1063 ASVs were retained across all 112 samples in the filtered dataset. Before analysing microbiome similarities across groups, we applied the variance stabilising transformation (VST) in DESeq2 (Love, Huber, & Anders, 2014), which uses a negative binomial mixed model to account for differences in library size across samples and to disentangle the relationship between the variance and the mean inherent to count data. Compared to other normalisation and transformation methods traditionally applied to microbiome data, the VST has the advantage of using all of the available data and is therefore preferable both to rarefying approaches (McMurdie & Holmes, 2014) and to transforming the data into relative abundances, which still has the problem of heteroscedasticity (Love et al., 2014). Based on the VS transformed data, we calculated Bray-Curtis dissimilarities (Bray & Curtis, 1957) among samples to visualise group differences using principle coordinate analysis (PCoA). We then statistically evaluated the microbiome composition in relation to sex, time point, host ID and health status using permutational multivariate analyses of variance (PERMANOVA, Anderson, 2001) with 1000 permutations using the adonis function in vegan (Oksanen et al., 2017). This approach is analogous to a parametric analysis of variance as that it partitions distance matrices into sources of variation and produces a pseudo-F value, the significance of which can be determined using a permutation test. As group differences detected using a PERMANOVA can be caused by variation in dispersion across groups rather than differences in mean values (Anderson, 2001), we tested for homogeneity of group dispersions using betadisper in vegan (Anderson, 2001; Oksanen et al., 2017) with *post-hoc* comparisons between specific contrasts evaluated with Tukey’s ‘honest significant differences’ method.

A main interest in microbial research is to determine the specific bacterial taxa that differ among groups. We therefore used the filtered but untransformed ASV data in combination with the DESeq2 method to determine differential abundances (Love et al., 2014). DESeq2 models abundance data such as microbial counts using a negative binomial distribution, estimates log fold changes between groups based on the specified model, and corrects the resulting *p*-values with a Benjamini and Hochberg false-discovery rate correction (Benjamini & Hochberg, 1995). As our ASV count matrix contained at least one zero in every row, we calculated the underlying size factors using the ‘poscounts’ estimator, which excludes zeros when calculating the geometric mean. To extract the appropriate group-specific contrasts, we fitted three different models and used a threshold of *p* < 0.01 to detect significant ASVs. Specifically, for analysing differential abundances between time points but within a given sex, the first two models contained ASV data for just females and just males respectively, while fitting both individual and time point in the model. To analyse and extract between-sex contrasts within each sampling time point, we constructed a third model by creating a new grouping factor as a combination of time point and sex, which was then fitted as predictor variable in the model.

To quantify which factors influence alpha diversity, we calculated Shannon indices based on the unfiltered and untransformed ASVs (2809 ASVs across 112 samples) to not bias the estimates by trimming rare ASVs, as suggested in the phyloseq tutorial (McMurdie & Holmes, 2013). Then we fitted a first Gaussian mixed model in lme4 (Bates, Mächler, Bolker, & Walker, 2014) with Shannon diversity as response, sex and time point as fixed effects and host ID as random effect. As we could only assess the health of the individuals at time point T1 and T3, during which we sampled blood, we fitted a second Gaussian mixed model including data from only these two time points with Shannon diversity as response, health status (healthy vs. not healthy), sex and time point as fixed effects and individual as random effect. We calculated the R^2^ based on (Nakagawa & Schielzeth, 2013) and 95% confidence intervals around the R^2^ and the model estimates using parametric bootstrapping with 1000 replications. The individual adjusted repeatability including 95% CI was estimated with rptR (Stoffel, Nakagawa, & Schielzeth, 2017) using the same model structure and 1000 bootstraps.

### Genetic relatedness and microbial similarity

We estimated pairwise genetic relatedness based on 21 microsatellite loci using the R package Demerelate (Kraemer & Gerlach, 2013). We used the Loiselle estimator (Loiselle, Sork, Nason, & Graham, 1995) which is unbiased for small sample sizes and converged towards stable values for the number of loci used in this study (Supplementary Figure 6). To match the microbial data to the pairwise genetic relatedness matrix containing 40 individuals for further analyses, we merged the microbial data across the three time points for every individual by summing up the ASV abundances. The 40 merged microbiome samples were then transformed using the variance-stabilising transformation in DEseq2 before calculating Bray-Curtis dissimilarities. Both the genetic relatedness matrix and the microbial dissimilarity matrix were then split by sex to calculate their correlation using a Mantel test implemented in the ecodist package (Goslee & Urban, 2007) using 10,000 bootstraps with the default resampling level of 0.9 to calculate confidence intervals and 10,000 permutations to test for statistical significance. We furthermore wanted to test for a difference in slopes between males and females, which is not possible using Mantel tests. Consequently, we fitted a simple linear model of microbiome similarity that included an interaction term between relatedness and sex. In this model, we essentially treated pairwise comparisons as data points, which makes the normal *p*-value meaningless due to pseudo-replication in the data. We therefore estimated the interaction slope and its confidence interval using parametric bootstrapping, and determined the corresponding *p*-value by randomly permuting the relatedness vector and re-fitting the model. This resulted in a distribution of interaction estimates and yielded the probability of seeing an effect as strong or stronger than the observed effect by chance.

We investigated the proportion of gut microbiota that are impacted by host genetics using a subsampling exercise. We began by calculating microbial similarities from the two most abundant ASVs and determining the strength of correlation of the resulting microbial similarity matrix with genetic relatedness. We then iteratively repeated this procedure while always adding the next two most abundant ASVs until we reached the complete dataset containing 1064 ASVs. Lastly, we wanted to know whether the correlation between genetic relatedness and microbial similarity changed across the three time points and if it differed between sexes. We therefore used the original unmerged dataset and subsetted both the microbial and genetic datasets six times to calculate and visualize the correlation for all three time points and both sexes.

## Results

To investigate the development of the gut microbiome in young northern elephant seals, we collected rectal swab samples from 40 animals during three time points after weaning. We started sampling immediately after their mothers stopped nursing and returned to the sea (time point T1) and then resampled each individual after two (T2) and four weeks (T3). The individuals were on average 28, 43 and 58 days old at time points T1, T2 and T3 respectively. As a few animals were lost or found dead during the study period (see Materials and methods for details), our final sample comprised a total of 112 rectal swabs across three time points for which we quantified bacterial communities using 16S rRNA sequencing. After assembling the raw reads into amplicon sequence variants (ASVs), we retained 1063 ASVs with an average of 286 ± 67 ASVs (mean ± sd) per sample.

### Characterization of the gut microbiome

Overall, the main bacterial phyla that we identified were typical of a mammalian gut microbiome (Figure 1), with the majority of ASVs belonging to the phyla Bacteroidetes (mean ± sd = 34% ± 2%), Firmicutes (mean ± sd = 29% ± 1%), Fusobacteria (mean ± sd = 19% ± 3%), and Proteobacteria (mean ± sd = 13% ± 1%). The relative abundances of these four phyla remained relatively stable over time, except for the Fusobacteria, which decreased steadily during weaning (Figure 1). However, at a finer taxonomic scale, we observed substantial changes across the three time points (see below).

**Figure 1:**
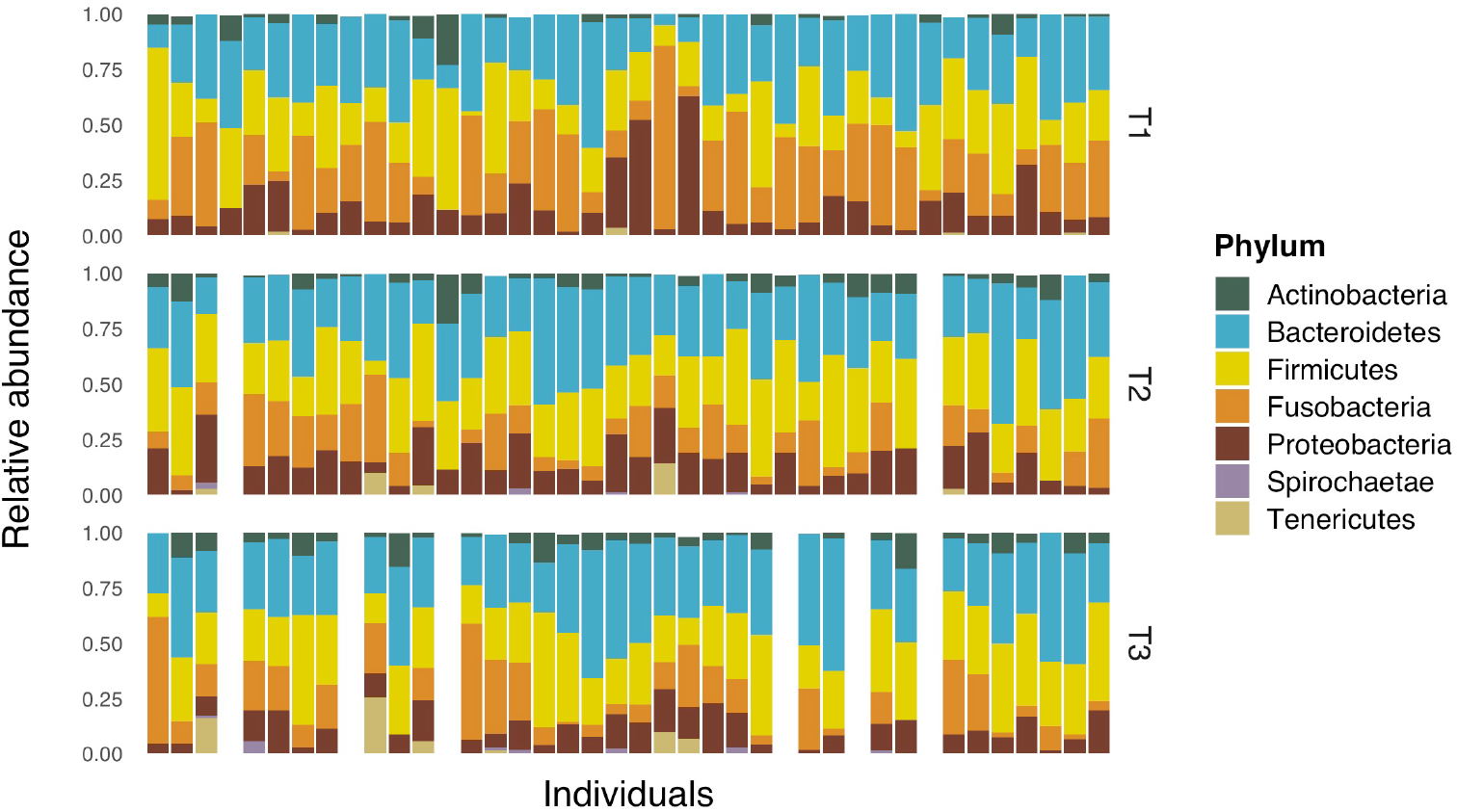
Relative abundance of bacterial phyla in gut samples from 40 northern elephant seals across three time points. The sampling time points T1, T2 and T3 correspond to individuals at 28, 43 and 58 days of age, respectively. Rare phyla with relative abundances below 1% are not shown. White columns represent individuals that either died or were lost during the course of the study.

### The core microbiome across individuals at different ages

We characterised the core microbiome at different developmental stages during the weaning period by extracting ASVs that appeared in at least 95% of samples at each time point (Supplementary Tables 1–3). Directly after weaning (T1), we identified 21 core ASVs, with two ASVs from the genera *Fusobacterium* and *Bacteroides* making up more than 25% of the average microbiome across individuals. This pattern changed substantially at T2 and T3. Here, we identified 15 and 35 core ASVs respectively, but the dominance of the two ASVs from T1 disappeared. Instead, a taxon from the genus *Ezakiella*, which only emerged after T1, became the most dominant ASV during T2 and T3 (with an average of 4% relative abundance). This is a recently discovered genus, of which only two species have been described; one from fecal samples of a coastal human indigeneous Peruvian population (Patel et al., 2015) and one from the human female genital tract (Diop, Raoult, Bretelle, & Fenollar, 2017). Closer to the time of nutritional independence (T3), a taxon from the genus *Prevotella* became the most successful new colonizer and the second most abundant genus. Concurrently, an ASV from the genus *Bacteroides*, which initially was the second most abundant taxon, decreased substantially in relative abundance (Supplementary Table 3, Supplementary Figure 4).

### Sex, age, host ID and health effects on gut microbiome beta diversity

To explore the major determinants of gut microbiome similarity across samples (beta diversity), we used a multidimensional scaling plot (MDS) of Bray-Curtis similarities between bacterial samples for visualisation and PERMANOVA (Anderson, 2001) for statistical analysis. Figure 2A shows variation due to sex and sampling time point (i.e. the age of individuals). Along the first axis, which accounts for 28.5% of the multidimensional spread in the data, a visible transition is apparent from the moment of weaning (T1) to the last sampling point (T3), shortly before the seals depart to the sea for feeding on their own, with samples from T2 being intermediate. A strong separation is also visible along axis 2, which accounts for 13.4% of the variation and reveals substantial differences in gut microbiome composition between the two sexes across the three sampling time points. Furthermore, microbiome samples from the same host cluster together, showing intra-individual consistency of gut microbial communities across the weaning-period (Figure 2B). Lastly, Figure 2C shows no or very little visible clustering of gut microbiome samples within healthy and non-healthy individuals, respectively.

**Figure 2:**
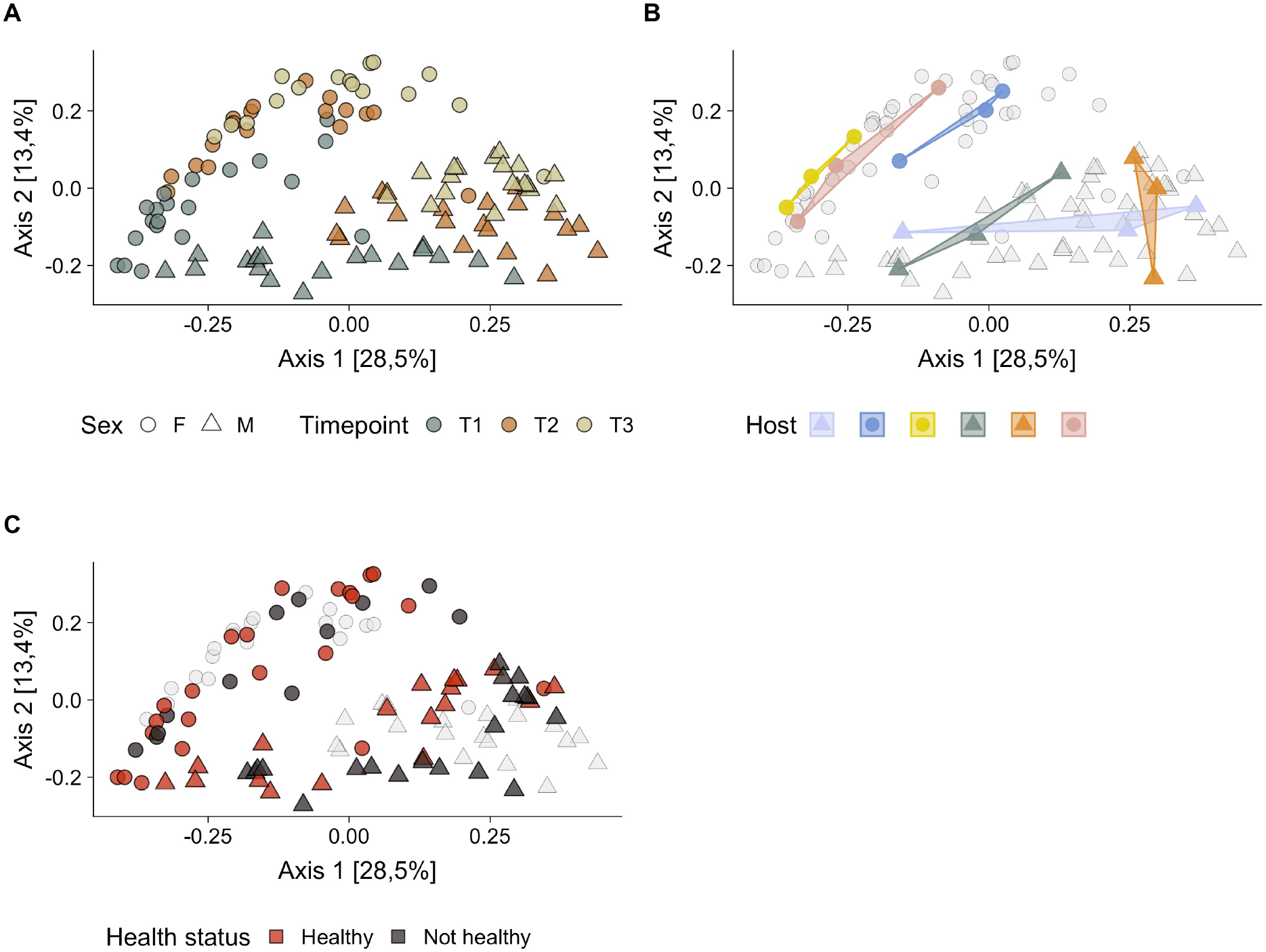
Gut microbiome sample beta diversity by (A) sex and sampling time point; (B) host ID and (C) health status. Shown are three different versions of the same multidimensional scaling (MDS) plot based on the Bray-Curtis similarities between 113 northern elephant seal gut microbiome samples, in which different color schemes are applied to emphasize different variables influencing beta diversity. All plots show samples obtained from males and females as rectangles and circles, respectively. Plot A is additionally colour coded according to sampling time point, plot B shows a selection of samples colour coded according to host identity (six samples were selected to avoid over-plotting while visualizing the similarity of microbiome samples obtained from the same host) and plot C shows colours according to the health status of the individuals. Health status could only be determined for samples at time point T1 and T3, when blood samples were taken. In all plots, females are denoted by circles and males by triangles. Data from all the samples were normalized using the variance stabilizing normalization implemented in DeSeq2 and the axes were length-scaled to reflect the Eigenvalues of the underlying principle coordinates.

To statistically analyse microbial group differences we fitted two PERMANOVA models which partition the microbial similarity matrix into variance components. The first model included sampling time point, sex and host ID as predictors of microbial similarity and included samples across all three time points. Overall, time point and sex each explain around 15% of the variation in microbial similarities (age: R^2^ = 0.15, *p* < 0.001, sex: R^2^ = 0.15, *p* < 0.001), while microbiome samples from the same host are also more similar, with host ID explaining 40% of the variation (R^2^ = 0.40, *p* < 0.001). After fitting the model with all of the samples, we analysed sex-differences within each time point *post-hoc* to avoid potential effects of repeated measures and to shed light on sex-specific patterns over time. The difference in gut microbiome between composition between males and females is already substantial at the time of weaning (T1: R^2^ = 0.13, *p* < 0.001), and further increases in the following weeks (T2: R^2^ = 0.26, *p* < 0.001, T3: R^2^ = 0.21, *p* < 0.001). Lastly, we compared specific time points *post-hoc*, while still fitting sex and host ID in the model. The transition from T1 to T2 explains 10.3% of the variation (R^2^ = 0.10, *p* < 0.001) while 4.1% is explained by microbial differences between T2 and T3 (R^2^ = 0.04, *p* < 0.001).

The second model included health status, sampling time point, sex and host ID as predictors of microbial similarity but was based only on samples from time points T1 and T3, during which we took blood samples to assess the health status of the individuals. Half of the pups sampled at T1 were clinically healthy, while most of the remaining pups had neutropenia (low levels of neutrophils) and lymphocytosis (high levels of lymphocytes), and 5% showed the opposite pattern, exhibiting neutrophilia and mild monocytosis. At T2, 39% of the pups had neutropenia and lymphocytosis, and only one pup had neutrophilia and mild monocytosis. Overall, health status only explained a negligible part of the overall variation in beta diversity (R^2^ = 0.025, *p* = 0.004). Consistent with the results from the first model, sex, time point and host ID each had large effects (age: R^2^ = 0.21, *p* < 0.001, sex: R^2^ = 0.11, *p* < 0.001, host ID: R^2^ = 0.44, *p* < 0.001). When comparing healthy and non-healthy individuals within each time point, the differences are slightly stronger at T1 (R^2^ = 0.06, *p* = 0.008) compared to T3 (R^2^ = 0.04, *p* = 0.116). The PERMANOVA assumption of multivariate homogeneity of group variances was met across all tests, as none of the contrasted groups differed in their dispersions (all *p* > 0.05). Consequently, all PERMANOVA results reflect differences in mean values across groups rather than differences in group dispersions (Anderson, 2001).

### Differential abundance of specific taxa across time and between sexes

At a finer scale, we used boxplots and raw data to visualize trends across time and sex for different hierarchical taxonomic ranks, from phylum to order. Supplementary Figures 1–3 reveal the complexity of the underlying dynamics, including multiple colonization and extinction events and often contrasting patterns at different taxonomic level. To quantify differences at the highest possible resolution, we tested for differentially abundant ASVs across time points and sexes using the DESeq2 method (Love et al., 2014). We provide a detailed description of all differential abundances in the Supplementary Material 2. Overall, the majority of significant changes in microbial abundances for both females and males happened between T1 and T2 (F: *n* = 100, M: *n* = 106) with less than half as many ASVs changing in abundance from T2 to T3 (F: *n* = 43, M: *n* = 26). Most of the ASVs that changed over time belonged to the *Clostridia* and *Bacteroidia* in both sexes (see Supplementary Figure 8). The number of differentially abundant ASVs between males and females was similarly large at all time points (T1: *n* = 96, T2: *n* = 102, T3: *n* = 80, see Supplementary Figure 9), and more than a third belonged to the *Clostridia Family XI* and the family *Ruminococcaceae*. However, while the overall number of differentially abundant ASVs between the sexes remained fairly similar throughout weaning, their taxonomic diversity appeared to increase (Supplementary Figure 9).

### Sex, age, host ID and health effects on gut microbiome alpha diversity

Microbial alpha diversity is frequently quantified in microbiome studies and is usually found to change during the development of vertebrates (Clark et al., 2015; O’Toole & Jeffery, 2015; Videvall et al., 2019). As a measure of alpha diversity, we used the Shannon index, which takes both species richness and the relative abundances of different species into account. To investigate the factors impacting microbial diversity across all three time points, we constructed a Gaussian mixed model of Shannon diversity with sex and sampling time point fitted as fixed effects and host ID as a random effect. The model only explained a small proportion of the variation in diversity (R^2^ = 0.06, 95% CI [0.01, 0.18]) but revealed a higher diversity for males than for females (β = 0.20, 95% CI [0.03, 0.39]). Moreover, Shannon diversity was stable across the post-weaning period and did not change between any two time points (T2 vs. T1: (β = 0.12, 95% CI [−0.10, 0.34], T3 vs. T1: β = 0.12, 95% CI [−0.07, 0.34]). These patterns are shown as boxplots alongside the raw data points in Figure 3. Contrary to microbial composition (beta diversity), where samples from the same host were more similar over time than between hosts, the alpha diversity of samples from the same individuals was not repeatable across time points (*r* = 0.1, 95% CI [0.00, 0.3]).

We then modelled the association between microbiome diversity and health status of individuals using a mixed model that only included data from the two time points in which we sampled blood and assessed the health of individuals. We fitted sex, health status and time point as fixed effects and host ID as a random effect. Healthy individuals hosted more diverse microbiomes than non-healthy individuals (β = 0.32, 95% CI [0.08, 0.55]), males had a slightly higher diversity than females (β = 0.15, 95% CI [−0.10, 0.40]) and the diversity was slightly higher at T3 than at T1 (β = 0.18, 95% CI [−0.04, 0.41]). Notably, this analysis suggests that the difference in diversity between males and females can be partially explained by a difference in the average health status of the two sexes, with the proportion of healthy individuals being higher in males (T1: F = 37%, M = 60%, T3: F = 33%, M = 44 %). Overall, this model explained slightly more variation in alpha diversity (R^2^ = 0.14, 95% CI [0.05, 0.36])

**Figure 3:**
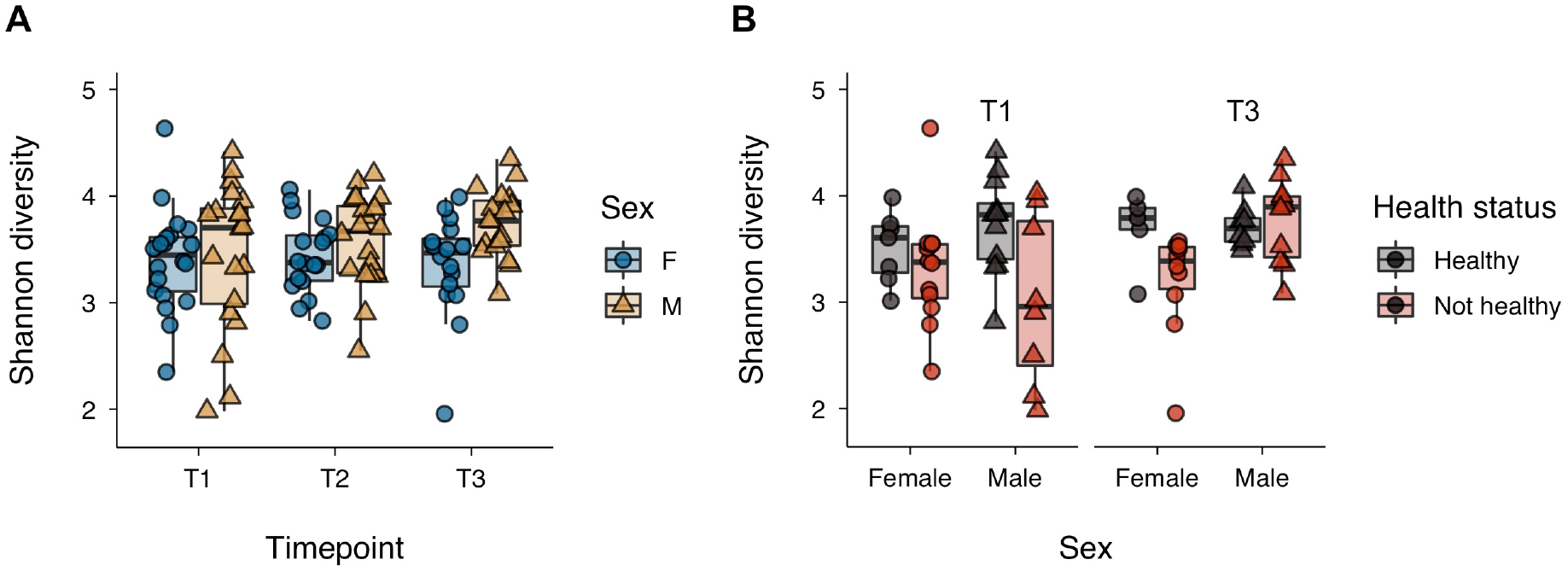
Sex-specific microbial alpha diversity (A) over time and (B) for healthy and non-healthy individuals within time points T1 and T3. Shown is the Shannon diversity of untransformed and unfiltered reads with circles and triangles representing samples from females and males, respectively. The boxplots are showing the intermediate 50% of data in the box and extending their whiskers to the data at maximally 1.5 times the interquartile range.

### Association between genetic relatedness and beta diversity

A fundamental topic in microbial ecology is the importance of host genotype for the formation of the gut microbiome. We approached this question by quantifying the correlation between host genetic relatedness and microbial similarity (Figure 4). Mantel tests showed a significant association in males (*r* = 0.26, CI [0.17, 0.34], *p* = 0.0013), which was visible across all three time points (Supplementary Figure 1). By contrast, we found no relationship in females, either overall (*r* = 0.06, CI [0.00, 0.12], *p* = 0.41) or within each time point (Supplementary Figure 4). As a difference in significance is not always a significant difference, we fitted a linear model to test for differences between the sex-specific slopes by fitting an interaction between relatedness and sex, with microbial dissimilarity as the response. The interaction term estimate was also negative (β = −0.11, CI [−0.23, −0.006], *p* = 0.02), indicating that microbial dissimilarity is more negatively correlated with genetic relatedness in males than in females.

**Figure 4:**
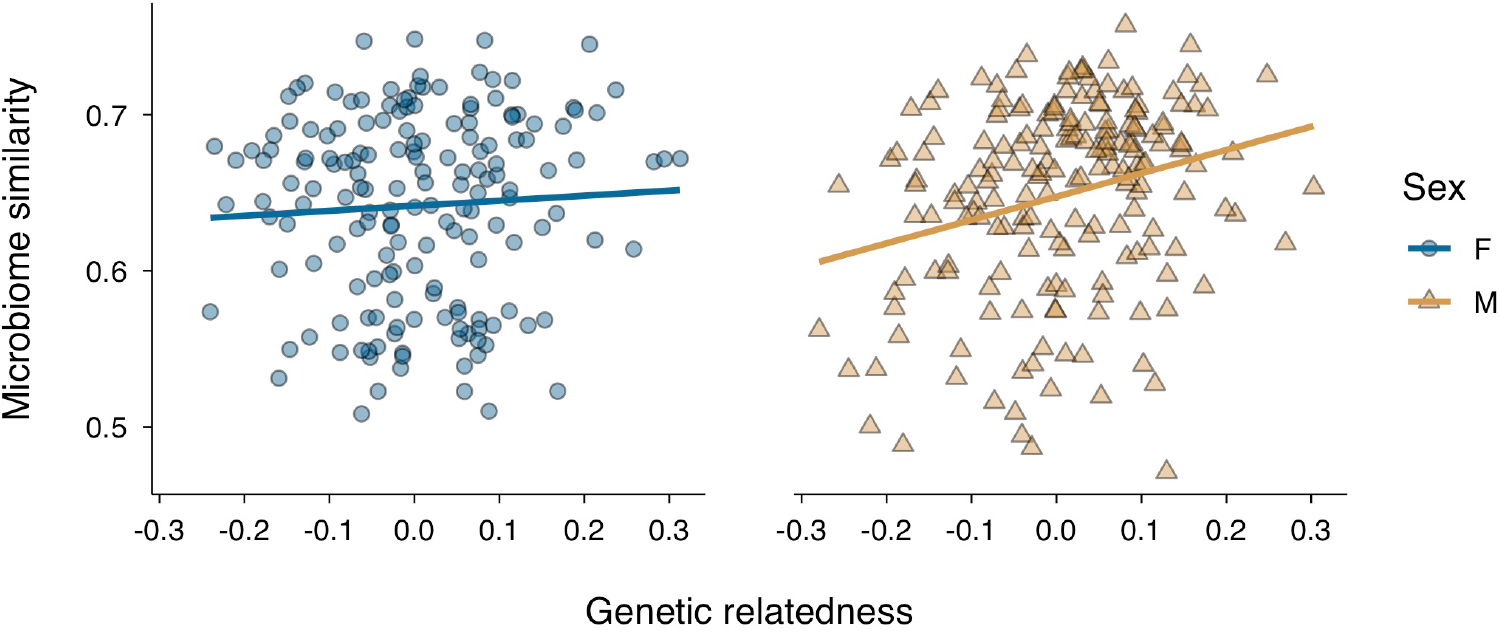
Relationship between pairwise gut microbiome similarity and genetic relatedness. For every individual, the microbiome data across all time points were merged by summing up ASV abundances. The abundance data were subsequently transformed using the variance-stabilising transformation in DEseq2 before calculating Bray-Curtis similarities between individuals. Genetic relatedness was estimated based on 21 microsatellite markers using the Loiselle estimator, with higher values representing elevated genetic relatedness.

To further investigate the effect of host relatedness on microbial similarity, we evaluated how many bacterial taxa are influenced by host genetics. We calculated the mantel correlation between genetic relatedness and microbial similarity based on an increasing number of ASVs, starting with the two most abundant (relative abundance) and iteratively increasing the number by the next two most abundant ASVs until we reached the full dataset (Figure 5). For females, the pattern across all subsets reflected the results from the full dataset and did not show a significant association between genetic relatedness and bacterial similarity. For males, a small number of ASVs contributed strongly to the overall correlation, but a peak in Mantel’s *r* was not reached until the 300 most abundant ASVs were included in the analysis. This suggests that a large proportion of taxa are at least slightly impacted by the host genotype as they contribute iteratively to an increasingly strong correlation.

**Figure 5:**
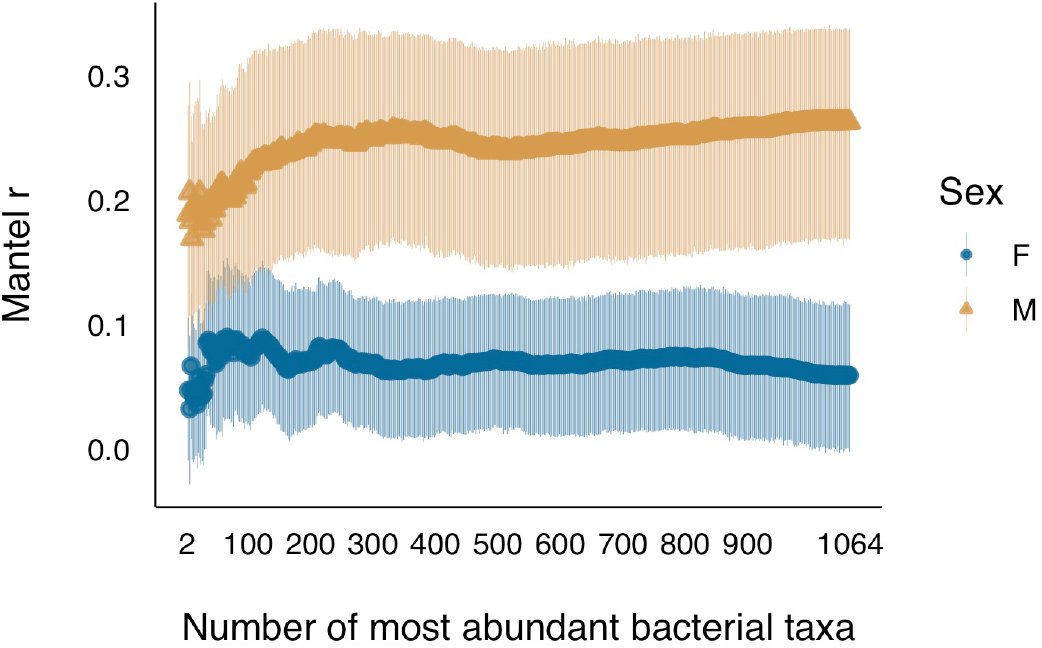
Strength of correlation between gut microbial similarity and genetic relatedness for an increasing number of microbial taxa. Each data point shows the correlation between microbial Bray-Curtis similarity and genetic relatedness with 95% confidence intervals calculated by non-parametric bootstrapping of samples. Bray-Curtis similarities were calculated based on an increasing number of ASVs, starting with the two ASVs yielding the highest relative abundances across all samples and iteratively increasing the number by the next two most abundant ASVs until the full dataset of 1064 ASVs was reached.

## Discussion

Microbiome studies of wild populations are essential for gaining an understanding of the ecological and evolutionary role of animal-microbe relationships (Hird, 2017). However, intrinsic effects on the gut microbiome such as sex differences are difficult to study, partly because they might be small in the first place, but also because environmental factors such as diet (David et al., 2014) may overshadow biological effects. Here, we studied an animal with extreme sexual dimorphism and sex-specific life-history strategies, where we would expect intra-species microbial variation to be large. Moreover, by sampling northern elephant seal weaners, which share the same dietary history and which did not feed during the sampling period, our sampling design minimized microbial variation due to diet, making it possible to provide a baseline assessment of the developing gut microbiome and shed light on individual-specific factors impacting it, in particular sex, age, health status and genotype.

First of all, we showed that the northern elephant seal gut microbiome at the time of weaning is already relatively complex with an average of nearly 300 ASVs per individual from 14 different phyla. Four of these phyla are highly abundant, the *Bacteroidetes*, *Firmicutes*, *Fusobacteria*, and *Proteobacteria*, and have previously been shown to be the main phyla in most pinniped guts (Bik et al., 2016; Glad et al., 2010; Nelson, Rogers, & Brown, 2013; Numberger, Herlemann, Jürgens, Dehnhardt, & Schulz-Vogt, 2016). However, their relative contribution varies across studies, partly due to differences across species, but also probably due to differences in sampling methods as well as environmental effects.

Patterns of early gut microbiome development are in general difficult to compare across mammals due to the small number of studies that have conducted longitudinal sampling during early life. One mammal in which the gut microbiome has been studied before and after weaning is the domestic pig, which shares some similarities to the patterns we report here. First of all, the most abundant bacterial phyla in weaned pigs, as in this study, were *Firmicutes* and *Bacteroidetes* (Pajarillo, Chae, Balolong, Kim, & Kang, 2014). Second, the dietary transition from nursing to weaning was found to be reflected in a marked decrease in the genus *Bacteroides* combined with an increase in *Prevotella* (Frese et al., 2015; Pajarillo et al., 2014), which we similarly observed in elephant seals. Although the specific functions of these genera are still under debate (Gorvitovskaia, Holmes, & Huse, 2016), *Bacteroides* have been shown to break down milk oligosaccharides and may therefore be important during nursing (A. Marcobal & Sonnenburg, 2012; Angela Marcobal et al., 2011), while *Prevotella* are associated with plant polysaccharide consumption and might therefore be important for the digestion of solid food (Ivarsson, Roos, Liu, & Lindberg, 2014). Interestingly, although increases in *Prevotella* have previously been associated with the transition to a solid diet (Frese et al., 2015), elephant seal weaners show an increase in *Prevotella* after weaning despite the fact that they are fasting. One possible explanation for this pattern could be a role of *Prevotella* for modulating immune tolerance instead of a diet-related function, which has previously been found in humans (Larsen, 2017).

Despite these changes in the composition of gut microbial communities, which include abundance changes as well as colonisation and extinction events, the average alpha diversity of elephant seal gut microbiomes was relatively stable throughout our study. This is surprising, as alpha diversity is usually quite dynamic across both shorter and longer time scales (Ballou et al., 2016; Frese et al., 2015; Kundu et al., 2017; Videvall et al., 2019). Consequently, the stability of gut microbial diversity observed in this study might be a consequence of the animals fasting during the study period, as dietary changes can be a major source of new microbial diversity (Pantoja-Feliciano et al., 2013).

In order to improve our understanding of host-microbe interactions in ecology and evolution, environmental sources of microbial variation need to be disentangled from individual-specific sources of variation. Sex is an important source of intraspecific differences, and hence a likely source of gut microbial variation. However, sex differences in the gut microbiome have mostly been found to be negligible or non-existent in wild populations so far (Bennett et al., 2016; Bobbie et al., 2017; Maurice et al., 2015; Ren et al., 2017; Tung et al., 2015). Although sex-differences may truly be small in some species, it is also possible that the effects of external factors such as diet or environment on gut microbial communities mask the effects of sex. In contrast to most of the literature, we found sex to be a strong and early determinant of gut microbiome composition, but not diversity, in elephant seals. In fact, sexual dimorphism in the gut microbiome composition precedes any morphological dimorphism, making it a precursor to what later becomes an extreme morphological and behavioural sexual dimorphism in adult elephant seals.

We found that apparent sex-differences in the gut microbiome alpha diversity (but not beta diversity) were largely explained by a higher proportion of healthy pups in males than in females. Animals that were categorized as healthy on the basis of their blood parameters possessed higher gut bacterial diversity than pups categorized as non-healthy. Recent evidence suggests that the gut microbiome impacts systemic immune effectors, including the development and output of circulating leukocytes (Grainger, Daw, & Wemyss, 2018) and other studies have shown that microbiome composition can reflect specific clinical conditions (Kozik, Nakatsu, Chun, & Jones-Hall, 2019; Pascal et al., 2017). Although none of the northern elephant seal pups we examined had evident signs of illness, and the observed deviations from normal blood values were mild, it is likely that the pups categorized as ‘non-healthy’ were suffering from some type of acute transient infection. Further investigation into associations between the gut microbiome, systemic and local immune effectors, and specific diseases could help to improve our understanding of links between clinical health and microbiome diversity. However, it is plausible that rather than signalling a direct relationship between microbiome diversity and health, our results could be a reflection of early sexual dimorphism in enteric immune tolerance, which would impact the establishment and stability of gut microbial communities (Duerr & Hornef, 2012; Fulde & Hornef, 2014).

A parallel can be drawn between our results and those of a study of southern elephant seals, which also detected sex specific differences in the gut microbiome (Nelson, Rogers, Carlini, & Brown, 2013). However, this particular study focused on adults, which show extreme sexual size dimorphism as well as marked sex specific differences in behaviour, diet and foraging behaviour, which would be expected to have strong effects on host associated microbial communities (Hindell et al., 2016). By contrast, male and female northern elephant seal pups are not visually distinguishable, and before our first sampling all pups remained close to their mothers to nurse, such that variation in behaviour, diet and social interactions was negligible. Furthermore, there is evidence for an equal maternal energy investment in female and male pups during nursing with respect to milk intake (Kretzmann, Costa, & Le Boeuf, 1993) and offspring mass change (Deutsch, Crocker, Costa, & Le Boeuf, 1994). It is therefore remarkable that males and females host very different gut microbiomes even directly after weaning, which could be due to early sex-specific intestinal adaptations and is consistent with earlier findings that diet can have sex-dependent effects on gut microbial communities (Bolnick et al., 2014). Whether sex-specific gut microbes are early signs of adaptation related to different adult feeding strategies or other life-history challenges will need to be examined in future studies.

A largely unanswered question is how strongly host genetics impacts the composition of the gut microbiome in wild populations. In humans and mice, genome-wide association studies have shown that at least a small proportion of the microbiome is genetically determined (Goodrich et al., 2016; Kurilshikov et al., 2017). It has also been shown that genetically more similar humans harbour more similar gut microbial communities (Zoetendal, Akkermans, Akkermans-van Vliet, de Visser, & de Vos, 2001), a pattern that seems difficult to replicate in natural populations (Degnan et al., 2012). In the wild, however, environmental factors such as diet, habitat or social behaviour are difficult to control for and are likely to mask any smaller, more subtle effects of host genotype. In this study, we found that genetically related males hosted more similar gut microbiomes. However, this was not the case in females, which also carry different gut microbiomes than males. This sex-specific relatedness-microbiome association is visible also within each sampling time point and is robust to the exclusion of a large number of microbial taxa. However, whether this difference is due to a temporal asymmetry in microbiome development among females and males or reflects more permanent sex-specific physiological mechanisms is still unclear.

## Conclusions

Northern elephant seals exhibit some of the most extreme sex differences among any mammal, both morphologically and in their life-histories. Here, we studied the gut microbiome and its development in northern elephant seal pups, from the time when their mothers abandoned them to shortly before they themselves head out to the open sea. Although morphologically speaking, male and female pups still cannot be distinguished apart during this period, we could show that the sexual dimorphism in their gut microbiome composition is already striking, a pattern that to our knowledge has not yet been found in any natural population and which is unlikely to be attributed to differences in diet. Within a natural, diet-controlled setting, we also showed that gut microbiome composition is associated with host genetic relatedness in males and changes substantially within only a few weeks after the end of lactation, potentially anticipating the growing elephant seals’ change in diet and life-style. Furthermore, we showed that health status has little impact on the beta diversity or composition of the microbiome although the alpha diversity is lower in clinically non-healthy individuals. We conclude that future gut microbiome studies of wild populations can benefit from species with large inter-individual variation such as northern elephant seals, and that minimising environmental variation and accounting for potential covariates will be crucial in order to gain a more in-depth understanding of microbial variation. Overall, our results provide a baseline assessment of the early colonization and development of elephant seal gut microbes, and contribute towards an improved understanding of host-microbe interactions in the wild, particularly in the light of sexual dimorphism.

## Acknowledgements

This research was supported by a Deutsche Forschungsgemeinschaft (DFG) standard grant (HO 5122/5-1) to JIH, and both a dual PhD studentship from Liverpool John Moores University and a DFG postdoctoral fellowship to MAS. The field work was funded by a Genetics Society Heredity fieldwork grant to MAS and by a grant from the Autonomous University of Queretaro (Fund for the Advancement of Scientific Research, 142908) awarded to KAW. The sequencing and laboratory work was also partially funded by the Animal Behaviour Department at Bielefeld University. We thank the Mexican Secretariat of the Environment and Natural Resources for the research permit, as well the Biosphere Reserve of the Pacific Islands off the Baja California Peninsula (CONANP, National Commission for Protected Areas).

## Data and code availability

The complete documented analysis pipeline and data to reproduce all analyses in the paper can be accessed via GitHub (https://github.com/mastoffel/nes_microbiome). The raw 16S sequences will be deposited on Dryad upon publication.

## Authors’ contributions

MAS, KAW and JIH conceived the study. MAS, KAW, NMD and FEV conducted the field work. MAS, SG and NC analysed the data. NC performed the genotyping. NMD conducted all haematology analyses. MAS, KAW and JIH wrote the paper and all authors gave feedback and comments.

