## supplementary_material for "Early sexual dimorphism in the developing gut microbiome of northern elephant seals"

Authors names and addresses:

Stoffel, M.A.[1,2,3*], Acevedo-Whitehouse, K.[4,5+], Nami Morales-Durán[4], Grosser, S.[6], Chakarov. N.[2], Krüger, O.[2], Nichols, H.J. [2,7], Elorriaga-Verplancken, F.R. [8], Hoffman, J.I.[2,9+]

[1]Institute of Evolutionary Biology, University of Edinburgh, Edinburgh, EH9 3FL, United Kingdom

[2]Department of Animal Behaviour, Bielefeld University, Postfach 100131, 33501 Bielefeld, Germany

[3]School of Natural Sciences and Psychology, Faculty of Science, Liverpool John Moores University, Liverpool L3 3AF, United Kingdom

[4]Unit for Basic and Applied Microbiology, School of Natural Sciences, Autonomous University of Queretaro, Avenida de las Ciencias S/N, Queretaro 76230, México

[5]The Marine Mammal Center. 2000 Bunker Road, Sausalito, CA 94965, USA.

[6]Division of Evolutionary Biology, Faculty of Biology, LMU Munich, Planegg-Martinsried, Germany

[7] Swansea University, Department of Biosciences, College of Science, Swansea University SA2 8PP, UK

[8]Department of Fisheries and Marine Biology, Centro Interdisciplinario de Ciencias Marinas, Instituto Politécnico Nacional (CICIMAR-IPN), La Paz, Mexico.

[9]British Antarctic Survey, High Cross, Cambridge, UK

Key index words:
gut microbiome, development, sex differences, life-history, northern elephant seal, wild

[+] Shared senior authors
* Corresponding author:
Martin A. Stoffel
Postal address: Institute of Evolutionary Biology, University of Edinburgh, Edinburgh, UK

### Supplementary Material 1 – Tables and Figures

| Kingdom | Phylum | Class | Order | Family | Genus | Species | Mean rel. abundance % |
|---|---|---|---|---|---|---|---|
| Bacteria | Fusobacteria | Fusobacteriia | Fusobacteriales | Fusobacteriaceae | Fusobacterium | NA | 17.22 |
| Bacteria | Bacteroidetes | Bacteroidia | Bacteroidales | Bacteroidaceae | Bacteroides | NA | 7.84 |
| Bacteria | Fusobacteria | Fusobacteriia | Fusobacteriales | Fusobacteriaceae | Fusobacterium | mortiferum | 2.04 |
| Bacteria | Fusobacteria | Fusobacteriia | Fusobacteriales | Fusobacteriaceae | Fusobacterium | mortiferum | 1.99 |
| Bacteria | Bacteroidetes | Bacteroidia | Bacteroidales | Bacteroidaceae | Bacteroides | fragilis | 1.67 |
| Bacteria | Fusobacteria | Fusobacteriia | Fusobacteriales | Fusobacteriaceae | Fusobacterium | NA | 1.22 |
| Bacteria | Firmicutes | Clostridia | Clostridiales | Family_XI | Anaerococcus | NA | 1.04 |
| Bacteria | Firmicutes | Clostridia | Clostridiales | Ruminococcaceae | NA | NA | 0.97 |
| Bacteria | Bacteroidetes | Bacteroidia | Bacteroidales | Porphyromonadaceae | Odoribacter | NA | 0.93 |
| Bacteria | Proteobacteria | Gammaproteobacteria | Aeromonadales | Succinivibrionaceae | Anaerobiospirillum | NA | 0.89 |
| Bacteria | Proteobacteria | Gammaproteobacteria | Enterobacteriales | Enterobacteriaceae | Escherichia/Shigella | NA | 0.72 |
| Bacteria | Proteobacteria | Gammaproteobacteria | Aeromonadales | Succinivibrionaceae | Anaerobiospirillum | NA | 0.60 |
| Bacteria | Firmicutes | Negativicutes | Selenomonadales | Acidaminococcaceae | Phascolarctobacterium | NA | 0.51 |
| Bacteria | Firmicutes | Clostridia | Clostridiales | Family_XI | Peptoniphilus | NA | 0.49 |
| Bacteria | Firmicutes | Clostridia | Clostridiales | Peptostreptococcaceae | Peptoclostridium | NA | 0.49 |
| Bacteria | Fusobacteria | Fusobacteriia | Fusobacteriales | Fusobacteriaceae | Fusobacterium | NA | 0.48 |
| Bacteria | Firmicutes | Negativicutes | Selenomonadales | Veillonellaceae | Dialister | NA | 0.44 |
| Bacteria | Firmicutes | Clostridia | Clostridiales | Ruminococcaceae | Anaerotruncus | NA | 0.35 |
| Bacteria | Firmicutes | Clostridia | Clostridiales | Lachnospiraceae | Blautia | NA | 0.14 |
| Bacteria | Actinobacteria | Coriobacteriia | Coriobacteriales | Coriobacteriaceae | Collinsella | NA | 0.11 |
| Bacteria | Bacteroidetes | Bacteroidia | Bacteroidales | Porphyromonadaceae | Parabacteroides | merdae | 0.08 |

**Supplementary Table 1:** Core microbiome shared among at least 95 % of samples during sampling time point one (T1). Every row represents an ASV, and a full table including the exact sequences is provided as Supplementary data (core_microbiome_T1.txt). In some cases, a taxonomic level could not be assigned (NA). Shown is also the mean relative abundance of each core ASV across all samples at T1.

| Kingdom | Phylum | Class | Order | Family | Genus | Species | Mean rel. abundance % |
|---|---|---|---|---|---|---|---|
| Bacteria | Firmicutes | Clostridia | Clostridiales | Family_XI | Ezakiella | NA | 4.37 |
| Bacteria | Fusobacteria | Fusobacteriia | Fusobacteriales | Fusobacteriaceae | Fusobacterium | NA | 3.22 |
| Bacteria | Bacteroidetes | Bacteroidia | Bacteroidales | Bacteroidaceae | Bacteroides | NA | 2.75 |
| Bacteria | Bacteroidetes | Bacteroidia | Bacteroidales | Bacteroidaceae | Bacteroides | fragilis | 2.30 |
| Bacteria | Bacteroidetes | Bacteroidia | Bacteroidales | Porphyromonadaceae | Odoribacter | NA | 1.40 |
| Bacteria | Fusobacteria | Fusobacteriia | Fusobacteriales | Fusobacteriaceae | Fusobacterium | mortiferum | 1.02 |
| Bacteria | Actinobacteria | Actinobacteria | Corynebacteriales | Corynebacteriaceae | Lawsonella | NA | 0.97 |
| Bacteria | Firmicutes | Negativicutes | Selenomonadales | Veillonellaceae | Dialister | NA | 0.94 |
| Bacteria | Firmicutes | Clostridia | Clostridiales | Family_XI | Peptoniphilus | NA | 0.86 |
| Bacteria | Fusobacteria | Fusobacteriia | Fusobacteriales | Fusobacteriaceae | Fusobacterium | mortiferum | 0.73 |
| Bacteria | Firmicutes | Clostridia | Clostridiales | Family_XI | Anaerococcus | NA | 0.69 |
| Bacteria | Firmicutes | Clostridia | Clostridiales | Ruminococcaceae | Ruminococcaceae_UCG-005 | NA | 0.50 |
| Bacteria | Firmicutes | Clostridia | Clostridiales | Family_XI | Anaerococcus | NA | 0.48 |
| Bacteria | Firmicutes | Clostridia | Clostridiales | Ruminococcaceae | Faecalibacterium | NA | 0.35 |
| Bacteria | Bacteroidetes | Bacteroidia | Bacteroidales | Rikenellaceae | Alistipes | NA | 0.24 |

**Supplementary Table 2:** Core microbiome shared among at least 95 % of samples during sampling time point two (T2). Every row represents an ASV, and a full table including the exact sequences is provided as Supplementary data (core_microbiome_T2.txt). In some cases, a taxonomic level could not be assigned (NA). Shown is also the mean relative abundance of each core ASV across all samples at T2

| Kingdom | Phylum | Class | Order | Family | Genus | Species | Mean rel. abundance % |
|---|---|---|---|---|---|---|---|
| Bacteria | Firmicutes | Clostridia | Clostridiales | Family_XI | Ezakiella | NA | 4.23 |
| Bacteria | Bacteroidetes | Bacteroidia | Bacteroidales | Prevotellaceae | Prevotella | NA | 4.22 |
| Bacteria | Bacteroidetes | Bacteroidia | Bacteroidales | Porphyromonadaceae | Porphyromonas | NA | 3.05 |
| Bacteria | Bacteroidetes | Bacteroidia | Bacteroidales | Porphyromonadaceae | Proteiniphilum | NA | 3.02 |
| Bacteria | Fusobacteria | Fusobacteriia | Fusobacteriales | Fusobacteriaceae | Fusobacterium | NA | 2.85 |
| Bacteria | Bacteroidetes | Bacteroidia | Bacteroidales | Bacteroidaceae | Bacteroides | NA | 2.68 |
| Bacteria | Bacteroidetes | Bacteroidia | Bacteroidales | Porphyromonadaceae | Porphyromonas | NA | 2.00 |
| Bacteria | Actinobacteria | Actinobacteria | Corynebacteriales | Corynebacteriaceae | Lawsonella | NA | 1.37 |
| Bacteria | Bacteroidetes | Bacteroidia | Bacteroidales | Bacteroidaceae | Bacteroides | NA | 1.11 |
| Bacteria | Firmicutes | Negativicutes | Selenomonadales | Veillonellaceae | Dialister | NA | 0.90 |
| Bacteria | Firmicutes | Clostridia | Clostridiales | Ruminococcaceae | Ruminococcaceae_UCG-005 | NA | 0.90 |
| Bacteria | Actinobacteria | Actinobacteria | Corynebacteriales | Corynebacteriaceae | Lawsonella | NA | 0.86 |
| Bacteria | Firmicutes | Clostridia | Clostridiales | Family_XI | Anaerococcus | NA | 0.86 |
| Bacteria | Firmicutes | Clostridia | Clostridiales | Family_XI | Anaerococcus | NA | 0.83 |
| Bacteria | Bacteroidetes | Bacteroidia | Bacteroidales | Porphyromonadaceae | Odoribacter | NA | 0.79 |
| Bacteria | Fusobacteria | Fusobacteriia | Fusobacteriales | Fusobacteriaceae | Fusobacterium | mortiferum | 0.75 |
| Bacteria | Firmicutes | Clostridia | Clostridiales | Ruminococcaceae | Ruminococcaceae_UCG-005 | NA | 0.71 |
| Bacteria | Fusobacteria | Fusobacteriia | Fusobacteriales | Fusobacteriaceae | Fusobacterium | mortiferum | 0.66 |
| Bacteria | Firmicutes | Clostridia | Clostridiales | Family_XI | Peptoniphilus | NA | 0.51 |
| Bacteria | Bacteroidetes | Bacteroidia | Bacteroidales | Bacteroidaceae | Bacteroides | fragilis | 0.48 |
| Bacteria | Firmicutes | Clostridia | Clostridiales | Ruminococcaceae | NA | NA | 0.46 |
| Bacteria | Firmicutes | Clostridia | Clostridiales | Ruminococcaceae | Anaerotruncus | NA | 0.43 |
| Bacteria | Firmicutes | Clostridia | Clostridiales | Ruminococcaceae | Ruminococcaceae_UCG-005 | NA | 0.40 |
| Bacteria | Firmicutes | Clostridia | Clostridiales | Ruminococcaceae | Ruminococcaceae_UCG-005 | NA | 0.30 |
| Bacteria | Fusobacteria | Fusobacteriia | Fusobacteriales | Fusobacteriaceae | Fusobacterium | NA | 0.28 |
| Bacteria | Firmicutes | Erysipelotrichia | Erysipelotrichales | Erysipelotrichaceae | NA | NA | 0.27 |
| Bacteria | Bacteroidetes | Bacteroidia | Bacteroidales | Rikenellaceae | Alistipes | NA | 0.14 |
| Bacteria | Firmicutes | Clostridia | Clostridiales | Ruminococcaceae | Anaerotruncus | NA | 0.11 |
| Bacteria | Firmicutes | Clostridia | Clostridiales | Ruminococcaceae | Ruminococcaceae_UCG-005 | NA | 0.11 |
| Bacteria | Firmicutes | Clostridia | Clostridiales | Family_XIII | NA | NA | 0.10 |
| Bacteria | Firmicutes | Clostridia | Clostridiales | Family_XI | Peptoniphilus | NA | 0.10 |
| Bacteria | Firmicutes | Clostridia | Clostridiales | Lachnospiraceae | Blautia | NA | 0.10 |
| Bacteria | Proteobacteria | Gammaproteobacteria | Pseudomonadales | Moraxellaceae | Psychrobacter | NA | 0.09 |
| Bacteria | Proteobacteria | Gammaproteobacteria | Pseudomonadales | Moraxellaceae | Psychrobacter | NA | 0.09 |
| Bacteria | Fusobacteria | Fusobacteriia | Fusobacteriales | Fusobacteriaceae | Fusobacterium | NA | 0.07 |

**Supplementary Table 3:** Core microbiome shared among at least 95 % of samples during sampling time point three (T3). Every row represents an ASV, and a full table including the exact sequences is provided as Supplementary data (core_microbiome_T3.txt). In some cases, a taxonomic level could not be assigned (NA). Shown is also the mean relative abundance of each core ASV across all samples at T3.

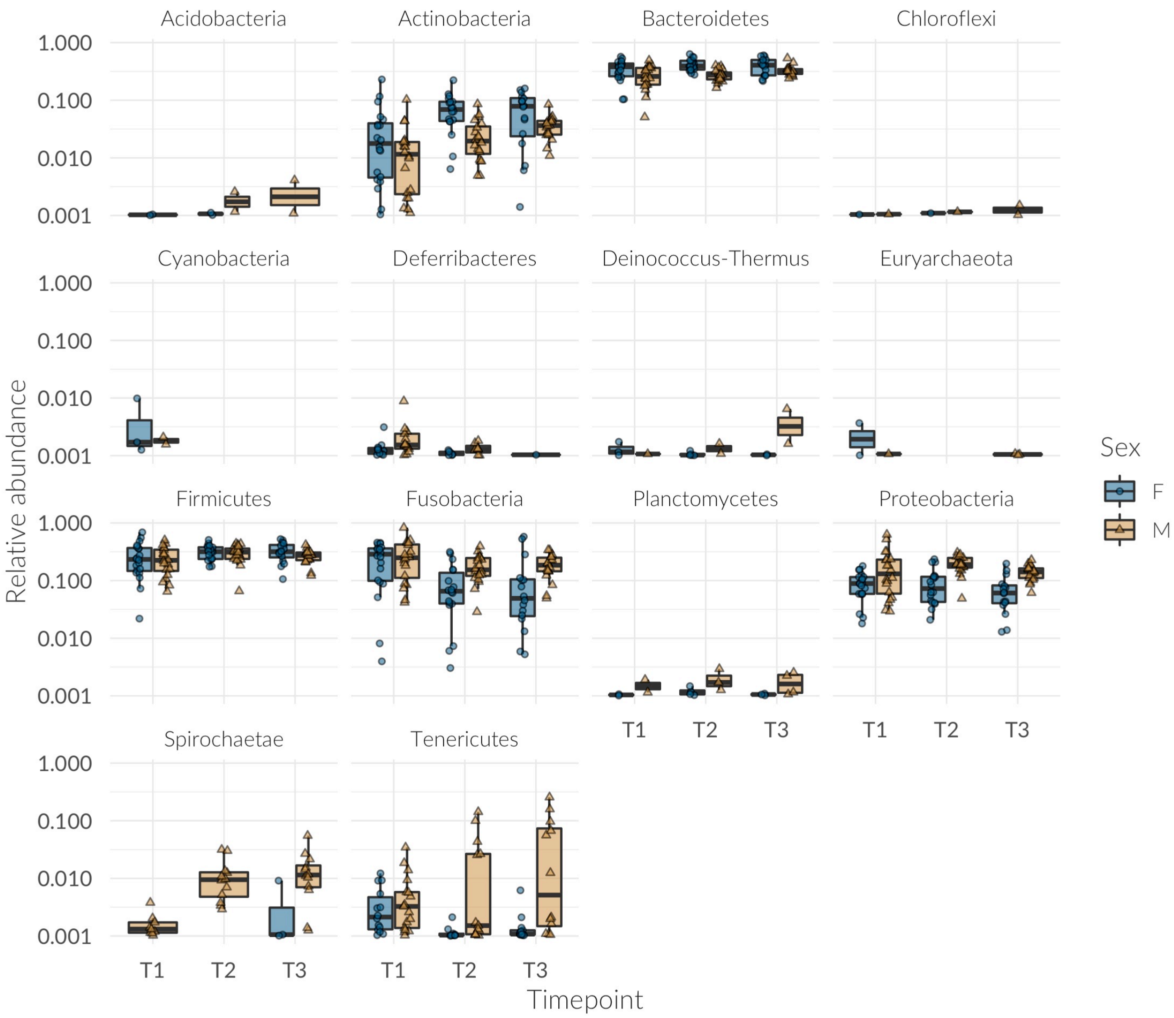


**Supplementary Figure 1:** Relative abundance of all bacterial taxa analysed in this study at the *Phylum* level across time points and colored by sex. Before visualization on the log scale, taxa with zero abundance were discarded and 0.001 added to all remaining relative abundances.

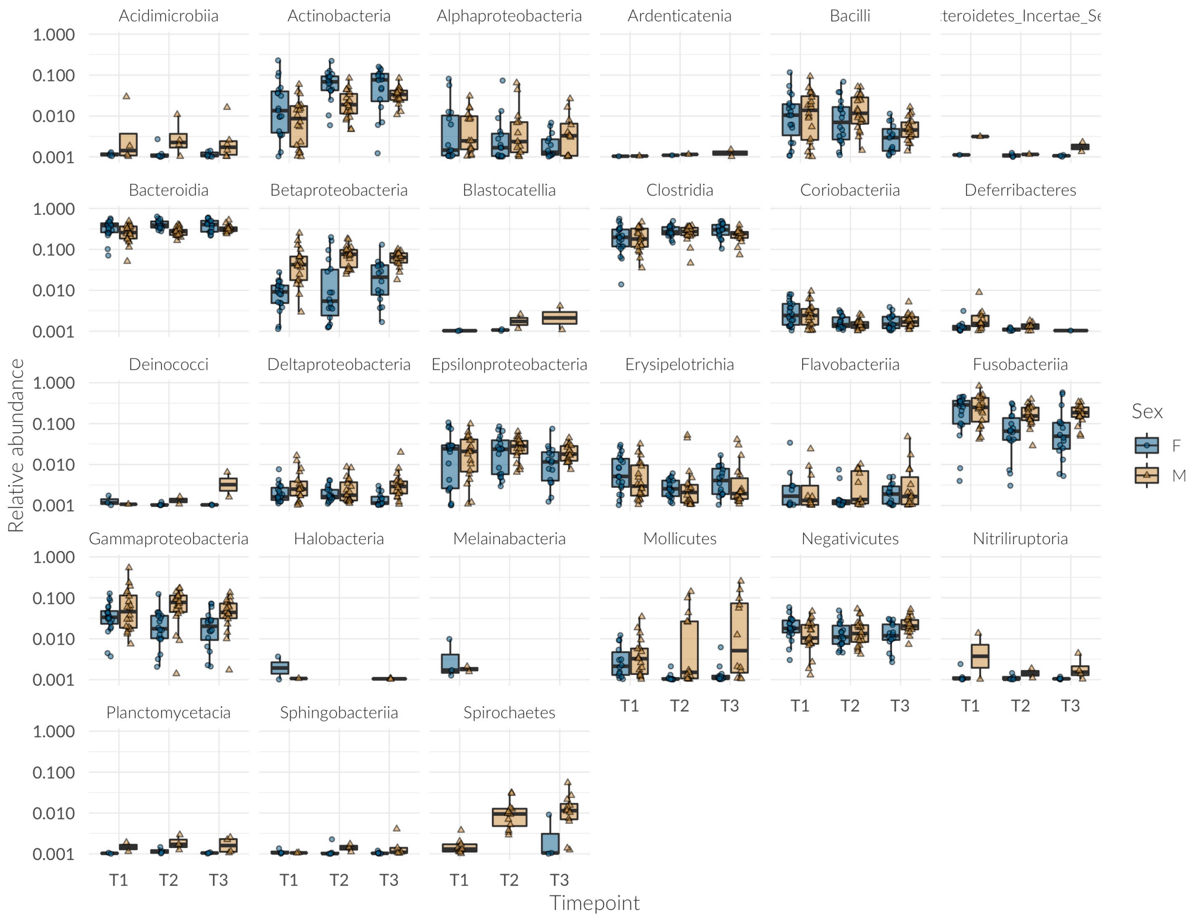


**Supplementary Figure 2**: Relative abundance of all bacterial taxa analysed in this study at the *Class* level across time points and colored by sex. Before visualization on the log scale, taxa with zero abundance were discarded and 0.001 added to all remaining relative abundances.

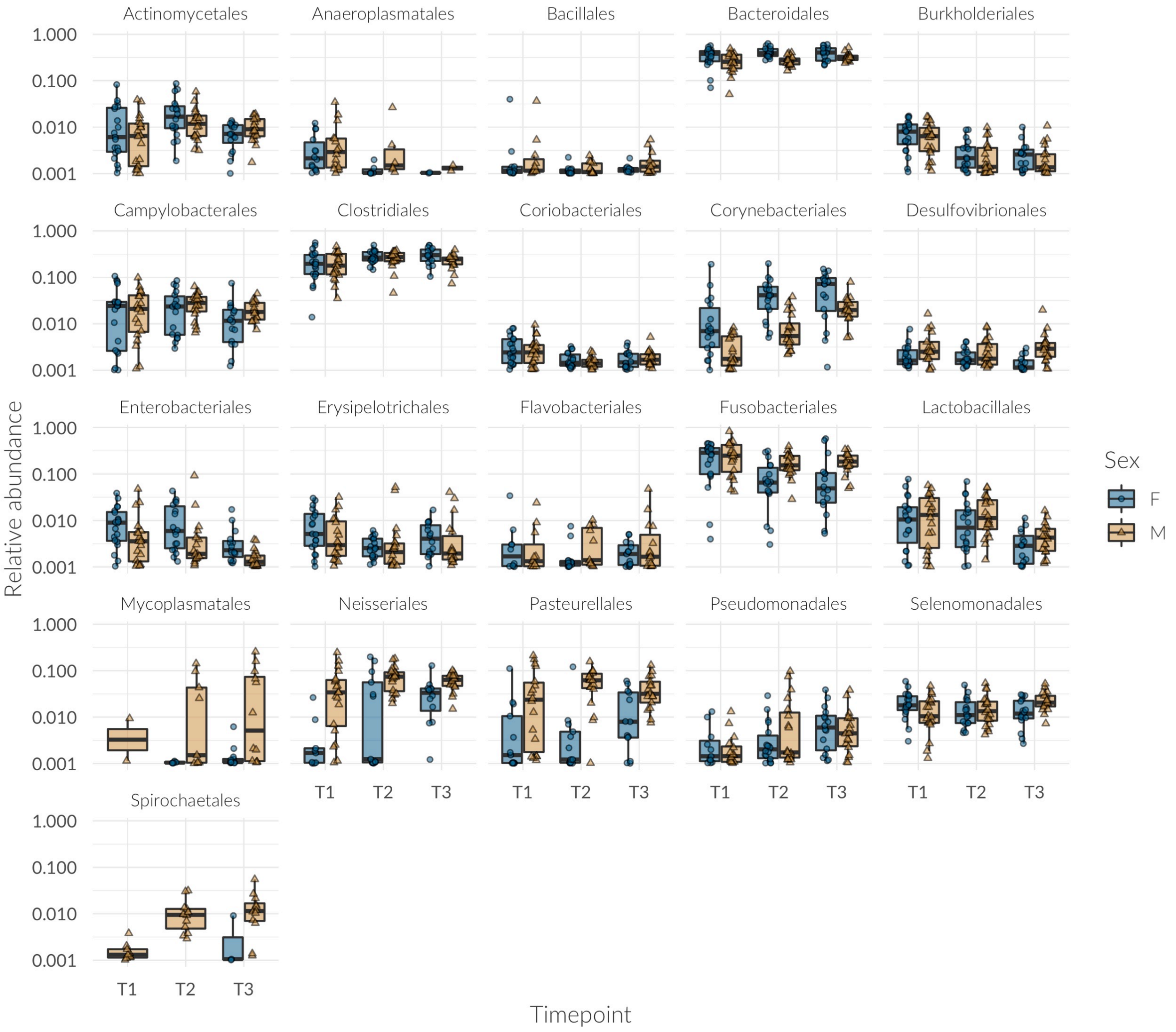


**Supplementary Figure 3:** Relative abundance of bacterial taxa analysed in this study at the *Order* level across time and colored by sex. Before visualization on the log scale, taxa with zero abundance were discarded and 0.001 added to all remaining relative abundances. Shown is a subset of bacterial orders with interesting patterns and/or high prevalence across samples.

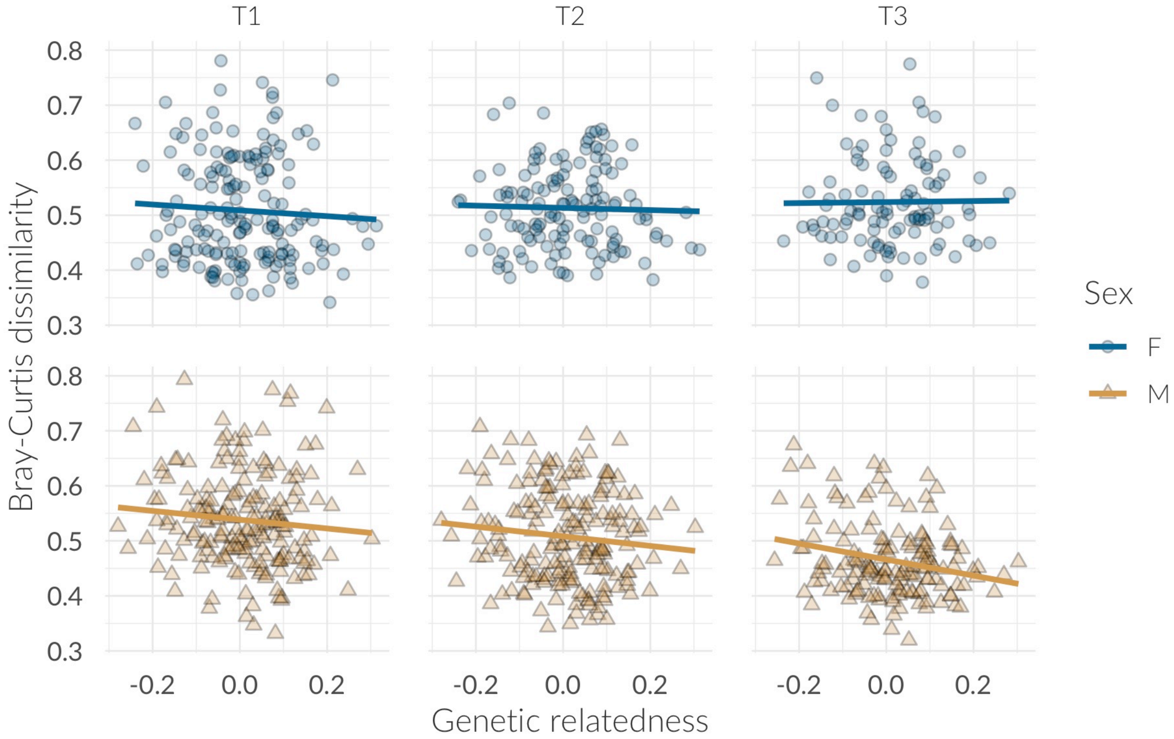


**Supplementary Figure 4:** Correlations between microbial similarity and genetic relatedness at three time points, split by sex.

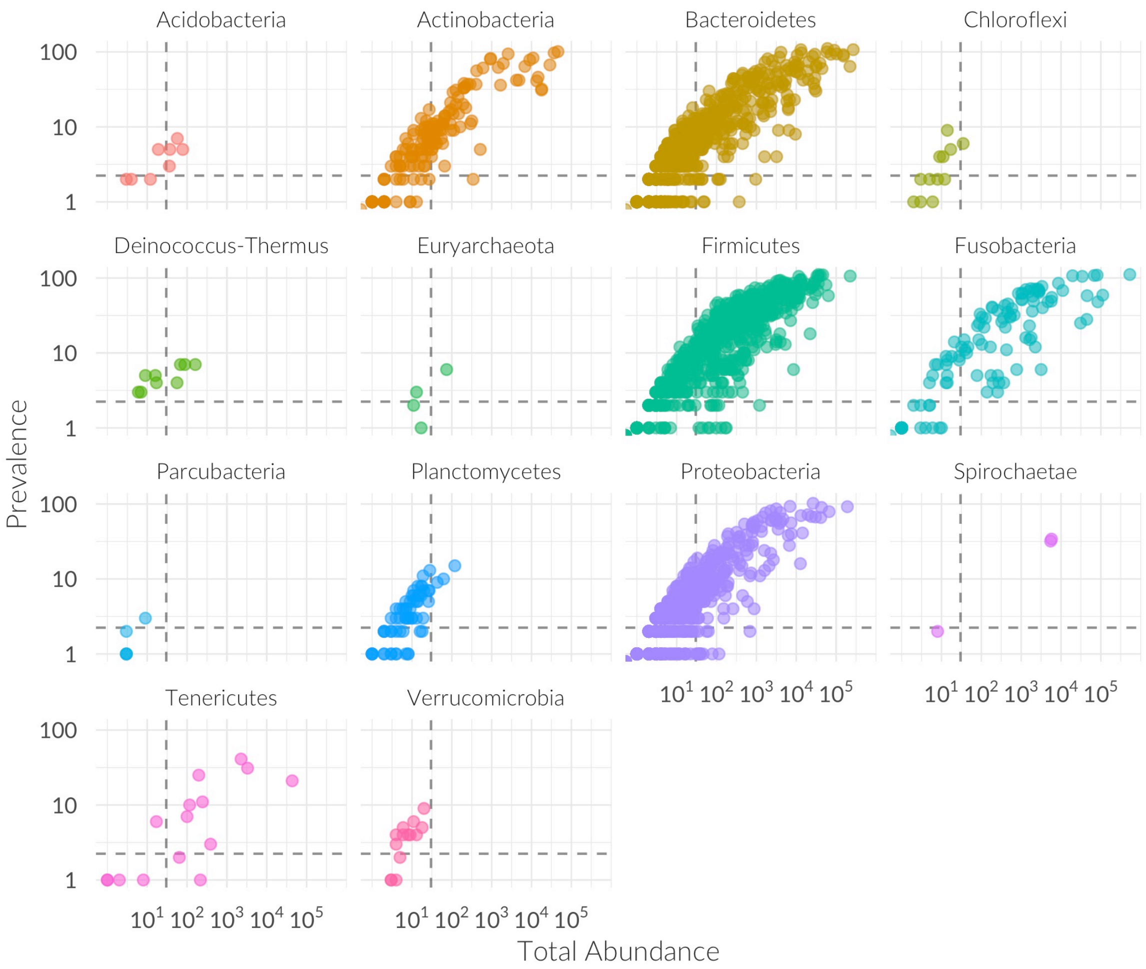


**Supplementary Figure 5**: Prevalence and total abundance of taxa split by phylum. The horizontal and vertical dashed line represent the cut-offs for filtering, with taxa present in fewer than three individuals and/or with an overall read count lower than 30 being discarded.

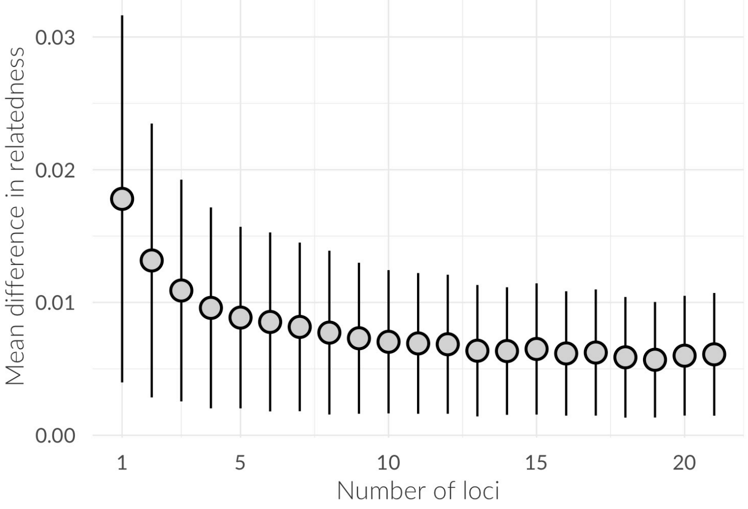


**Supplementary Figure 1: Sensitivity of the Loiselle relatedness estimator to the number of loci used.** Plotted are the mean and standard deviation (SD) of differences in pairwise genetic relatedness against the number of loci used. SDs were calculated from 1000 bootstrap replicates per locus number.

### Supplementary Material 2 – Differential abundances of specific taxa

**Differential abundance of specific taxa with age**

Despite the apparent similarity of phyla across all three time points (Figure 1), on a finer scale a large number of bacterial taxa changed in abundance over time (Supplementary Figure 7 and 8). Most significant changes happen early on, with a large number of taxa for each sex varying from T1 to T2 (F: n = 100, M: n = 106) followed by a smaller number of significantly different abundances of taxa between T2 and T3 (F: n = 43, M: n = 26). On a taxonomic scale, most bacterial classes change substantially (Supplementary Figure 7). Between T1 and T2 most of the bacteria that change abundance belong to the Clostridia in both sexes (F: 47%, M: 44%), followed by Bacteroidia (F: 18%, M: 20%) and Fusobacteria (F:13%, M: 12%), a pattern which is very similar for the second transition between T2 and T3 in males (Clostridia 35%, Bacteriodia 19%, Fusobacteria 15%) while in females the Bacteriodia (37%) change most substantially, more so than the Clostridia (30%) and Gammaproteobacteria (14 %). Several interesting changes also happen in the less abundant bacterial classes. While Deferribacteres go extinct over time, the Spirochaetes increase in abundance mainly in males (Supplementary Figure 3) and start to colonise females at T3. The Bacilli and the Fusobacteria deplete quickly over time, while the Actinobacteria increase in their relative abundances by nearly ten-fold in females and by more than five-fold in males (Supplementary Figure 2).

**Sex specific patterns of changes**

Bacterial communities in both sexes show similar dynamics throughout the weaning period, although the 'baseline' abundances of many species differ substantially (Supplementary Figures 1-3, Supplementary Figure 8). On the phylum level, the microbial shift from T1 to T2 in both females and males consists mostly of taxa belonging to the Firmicutes (F: 51%, M: 48%), followed by Bacteroidetes (F: 18%, M: 20%) and Fusobacteria in males (13%) but Proteobacteria in females (14 %). Interestingly, a few bacterial families undergo large changes in abundance from T1 to T2 and make up a major part of the significantly different taxa, especially the Ruminococcaceae (F: 22%, M: 19%), followed by Fusobacteriaceae (F: 12%, M: 10%) and Lachnospiraceae in females (12%) but Porphyromonadaceae in males (9%). Bacterial changes between T2 and T3 mainly occurred in the phyla Bacteroidetes (37%), Firmicutes (32%) and Proteobacteria (19%) in females and Firmicutes (46%), Bacteroidetes (19%), Fusobacteria (15%) and Proteobacteria (15%) in males. Most differentially abundant taxa belonged, similar to the first transition, to the Ruminococcaceae (F:12%, M:23%), Porphyromonadaceae (F:16%, M:12%), and the Lachnospiraceae (12%) in females as well as the Leptotrichiaceae (12%) in males.

**Differential abundance of taxa across the sexes**

Despite similar dynamics over time, many taxa were significantly differentially abundant in males and females within all three time points (T1: n = 96, T2: n = 102, T3: n = 80, see Figure 5 and Figure 3). Although many phylogenetically different taxa contribute to these sex differences, three families contributed disproportionally much. The Clostridiales Family XI contributed 15% of differentially abundant taxa at T1, 16% at T2, and 18% at T3. The Ruminococcaceae contributed 15% of the taxa at T1, 19 % at T2 and 13 % at T3. The Porphyromonadaceae make up large differences at T1 (13%) and T2 (12%) but less so at T3 (4%).

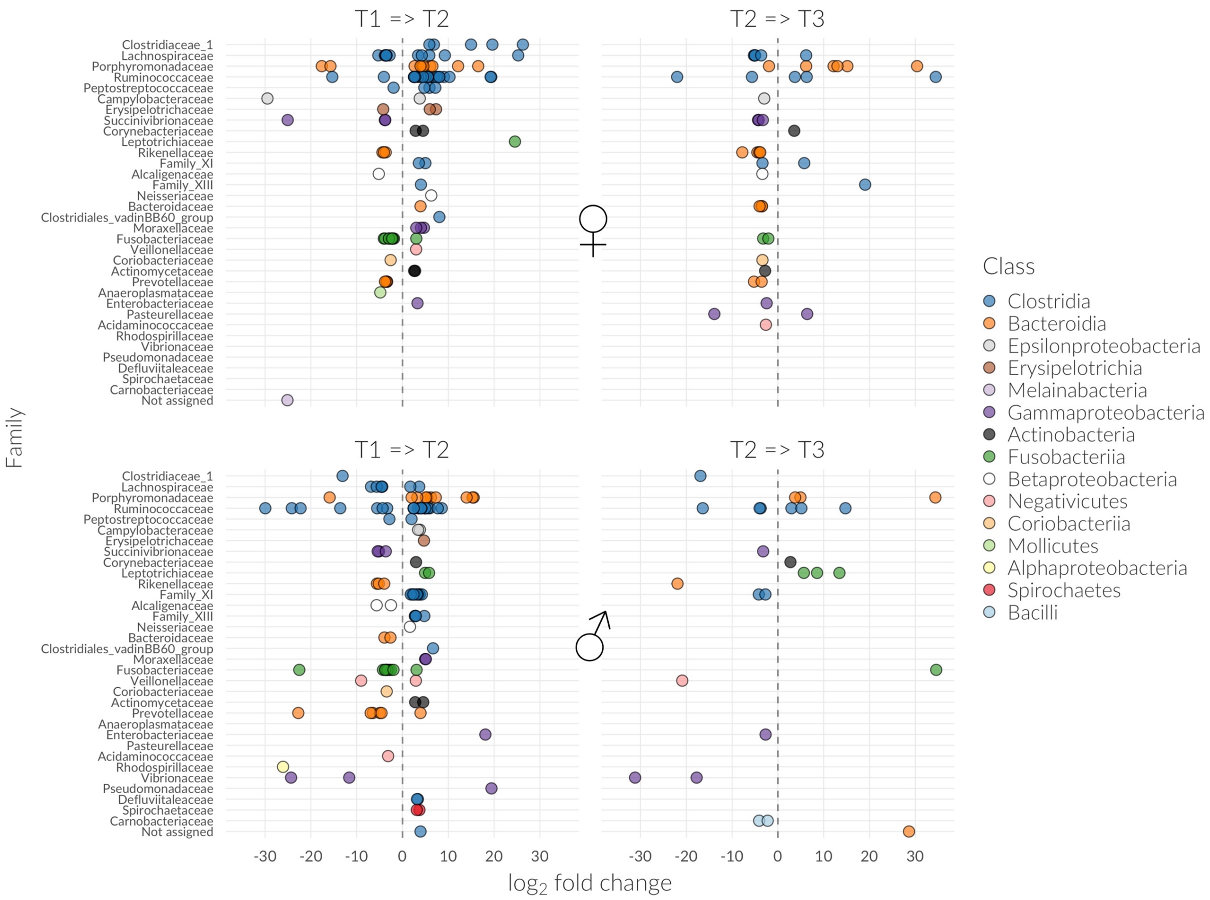


Supplementary Figure 7: Differential abundance of taxa between sampling points, split by sex.

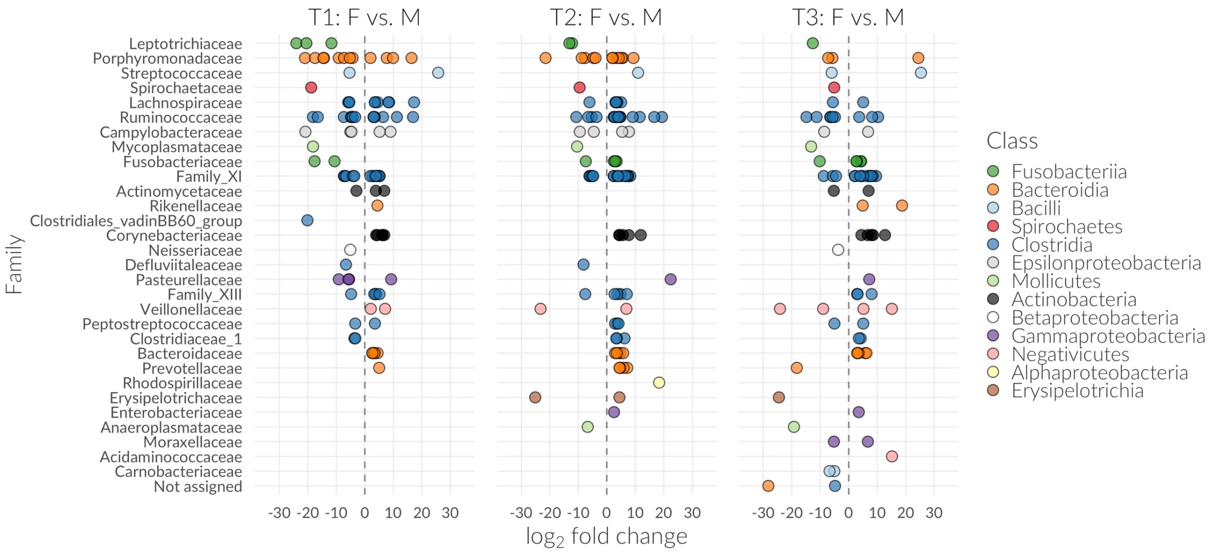


Supplementary Figure 8: Differential abundance of microbes between sex, split by sampling points

#### Supplementary Material 3 - Genotyping methods

Total genomic DNA of 40 *Mirounga angustirostris* samples was extracted from each sample using silica-gel membrane technology (DNeasy Blood and Tissue kit, Qiagen) and genotyped at 21 previously developed microsatellite loci (see Supplementary Table 4 for details). The microsatellite loci were amplified in singleplex or multiplex reactions. The following PCR profile was used: one cycle of 3 min at 94 °C; 30 cycles of 30 s at 94 °C, 30 s at Ta °C and 40 s at 72 °C; 8 cycles of 30 s at 94 °C, 30 s at 47 °C and 40 s at 72 °C; and one final cycle of 10 min at 72 °C (see Supplementary Table 14 for $T_a$). Magnesium concentrations varied among the PCR mastermixes as shown in Supplementary Table 14. Fluorescently labelled PCR products were resolved by electrophoresis on an ABI 3730xl capillary sequencer and allele sizes were scored automatically using GeneMarker v1.85. To ensure high genotype quality, all traces were manually inspected and any incorrect calls were adjusted accordingly.

| Locus | Literature source | Mg (mM) | $T_a$ (°C) |
|---|---|---|---|
| 71HDZ441 | Huebinger et al. (2007) | 1.5 | 54 |
| Hg4.2 | Allen et al. (1995) | 1.5 | 56 |
| Lw-8 | Davis et al. (2002) | 1.5 | 47 |
| ZcCgDh4.7 | Hernandez-Velazquez et al. (2005) | 1.75 | 56 |
| PV9 | Goodman (1997) | 2 | 53 |
| ZzCgDh3.6 | Hernandez-Velazquez et al. (2005) | 2 | 39 |
| Hl-8 | Davis et al. (2002) | 2 | 53 |
| PVC1 | Garza (1998) | 1.5 | 52 |
| 71HDZ301 | Huebinger et al. (2007) | 1.5 | 42 |
| ZzCgDh1.8 | Hernandez-Velazquez et al. (2005) | 1.5 | 42 |
| ZcwA12 | Hoffman et al. (2007) | 1.75 | 49 |
| ZcwF07 | Hoffman et al. (2007) | 1.75 | 49 |
| Ag-9 | Hoffman et al. (2008) | 2 | 57 |
| ZcwC01 | Hoffman et al. (2007) | 2 | 57 |
| ZcwE04 | Hoffman et al. (2007) | 2 | 52 |
| ZcwG04 | Hoffman et al. (2007) | 2 | 52 |
| Mango01 | (Sanvito et al., 2013) | 1.5 | 55 |
| Mango44 | (Sanvito et al., 2013) | 1.5 | 55 |
| Mango43 | (Sanvito et al., 2013) | 1.5 | 55 |
| Mango35 | (Sanvito et al., 2013) | 1.5 | 53 |
| Mango06 | (Sanvito et al., 2013) | 1.5 | 55 |
| Mango09E19 | (Sanvito et al., 2013) | 1.5 | 52 |
| PV9.1 | This study | 1.5 | 53 |

**Supplementary Table 4: Microsatellite loci genotyped in the northern elephant seal.** "Mg" denotes the concentration of magnesium used in the PCR mastermix and "$T_a$" denotes the annealing temperature used.
